# Mechanism of target site selection by type V-K CRISPR-associated transposases

**DOI:** 10.1101/2023.07.14.548620

**Authors:** Jerrin Thomas George, Christopher Acree, Jung-Un Park, Muwen Kong, Tanner Wiegand, Yanis Luca Pignot, Elizabeth H. Kellogg, Eric C. Greene, Samuel H. Sternberg

## Abstract

Unlike canonical CRISPR-Cas systems that rely on RNA-guided nucleases for target cleavage, CRISPR-associated transposases (CASTs) repurpose nuclease-deficient CRISPR effectors to facilitate RNA-guided transposition of large genetic payloads. Type V-K CASTs offer several potential upsides for genome engineering, due to their compact size, easy programmability, and unidirectional integration. However, these systems are substantially less accurate than type I-F CASTs, and the molecular basis for this difference has remained elusive. Here we reveal that type V-K CASTs undergo two distinct mobilization pathways with remarkably different specificities: RNA-dependent and RNA-independent transposition. Whereas RNA-dependent transposition relies on Cas12k for accurate target selection, RNA-independent integration events are untargeted and primarily driven by the local availability of TnsC filaments. The cryo-EM structure of the untargeted complex reveals a TnsB-TnsC-TniQ transpososome that encompasses two turns of a TnsC filament and otherwise resembles major architectural aspects of the Cas12k-containing transpososome. Using single-molecule experiments and genome-wide meta-analyses, we found that AT-rich sites are preferred substrates for untargeted transposition and that the TnsB transposase also imparts local specificity, which collectively determine the precise insertion site. Knowledge of these motifs allowed us to direct untargeted transposition events to specific hotspot regions of a plasmid. Finally, by exploiting TnsB’s preference for on-target integration and modulating the availability of TnsC, we suppressed RNA-independent transposition events and increased type V-K CAST specificity up to 98.1%, without compromising the efficiency of on-target integration. Collectively, our results reveal the importance of dissecting target site selection mechanisms and highlight new opportunities to leverage CAST systems for accurate, kilobase-scale genome engineering applications.

## INTRODUCTION

Bacteria encode diverse mobile genetic elements that exhibit a wide spectrum of mobilization behaviors, ranging from selective targeting of fixed attachment sites to promiscuous insertion into degenerate sequence motifs^1^. Although insertion specificity is often dictated by a single recombinase enzyme^2, 3^, some transposons encode heteromeric transposase complexes that distribute DNA target and DNA integration activities across multiple distinct molecular components^4, 5^. Tn*7*-like transposons are unique in this regard, in that they have evolved to exploit diverse molecular pathways for target site selection, including site-specific DNA binding proteins^4–6^, replication fork-specific DNA binding proteins^7–9^, CRISPR RNA-guided DNA binding complexes^10–12^, and additional DNA targeting pathways that have yet to be characterized^13^. CRISPR-associated transposons (CASTs), in particular, represent both a fascinating example of CRISPR-Cas exaptation as well as an opportune starting point for the development of next-generation tools for programmable, large-scale DNA insertion^14, 15^.

CAST systems characterized to date fall within either type I or type V classes, which differ in their reliance on either Cascade or Cas12k effector complexes, respectively^10–12, 16, 17^. Although the core transposition machinery is conserved across CAST families — and includes a DDE-family transposase for integration (TnsB), a AAA+ ATPase for target site selection (TnsC), and an adaptor protein for CRISPR-transposition coupling (TniQ) — key molecular features distinguish the integration behavior of archetypal type I-F and type V-K systems. Whereas second-strand cleavage is catalyzed by the TnsA endonuclease in type I-F CASTs, leading to cut-and-paste transposition products, type V-K CASTs lack TnsA and instead mobilize through a copy-and-paste process, yielding cointegrate products^18–20^. Type I-F CASTs achieve single-digit genomic integration efficiencies when expressed in mammalian cells, as opposed to low but detectable activity only on ectopic plasmid targets for an improved type V-K CAST homolog^20, 21^. Additionally, heterologous expression of the CAST machinery from both systems yields vastly different integration specificities, with VchCAST (I-F) exhibiting mostly on-target activity in bacterial cells, as compared to an abundance of off-target insertions catalyzed by ShCAST (V-K)^11, 14^. Despite these differences, it is to be noted that type V-K CASTs have a compact coding sequence composed of four components compared to type I-F CAST (1666 vs. 2748 amino acids) and integrates predominantly in a unidirectional orientation^11^. The molecular basis underlying these distinguishing properties remains unexplored, particularly for type V-K CAST systems, limiting their practical application.

Recent structural studies have provided novel insights into the overall architecture of RNA-guided, ShCAST transpososome complexes^22, 23^. Target sites are marked by Cas12k binding^24, 25^, in conjunction with TniQ and ribosomal protein S15, which engages the tracrRNA component^22^, leading to stable R-loop formation reminiscent of other CRISPR effectors. In a key next step that is still poorly understood, TnsC assembles into filaments around double-stranded DNA — which can form adjacent to bound Cas12k-TniQ complexes^22^ or on naked DNA^24, 26^ — acting as a platform for the subsequent recruitment of the TnsB transposase that is scaffolded along conserved binding sites in the transposon left and right ends. DNA integration then occurs via a concerted transesterification at sites exposed by the TnsC filament, leading to transposons inserted at a fixed spacing downstream of the Cas12k-bound target site^11, 23^. Whether a similar assembly pathway is operational at the many off-target integration events observed with ShCAST expression in cells, or whether these represent an alternative transposition pathway, has not been systematically explored (**Fig. 1a**).

**Fig. 1.**
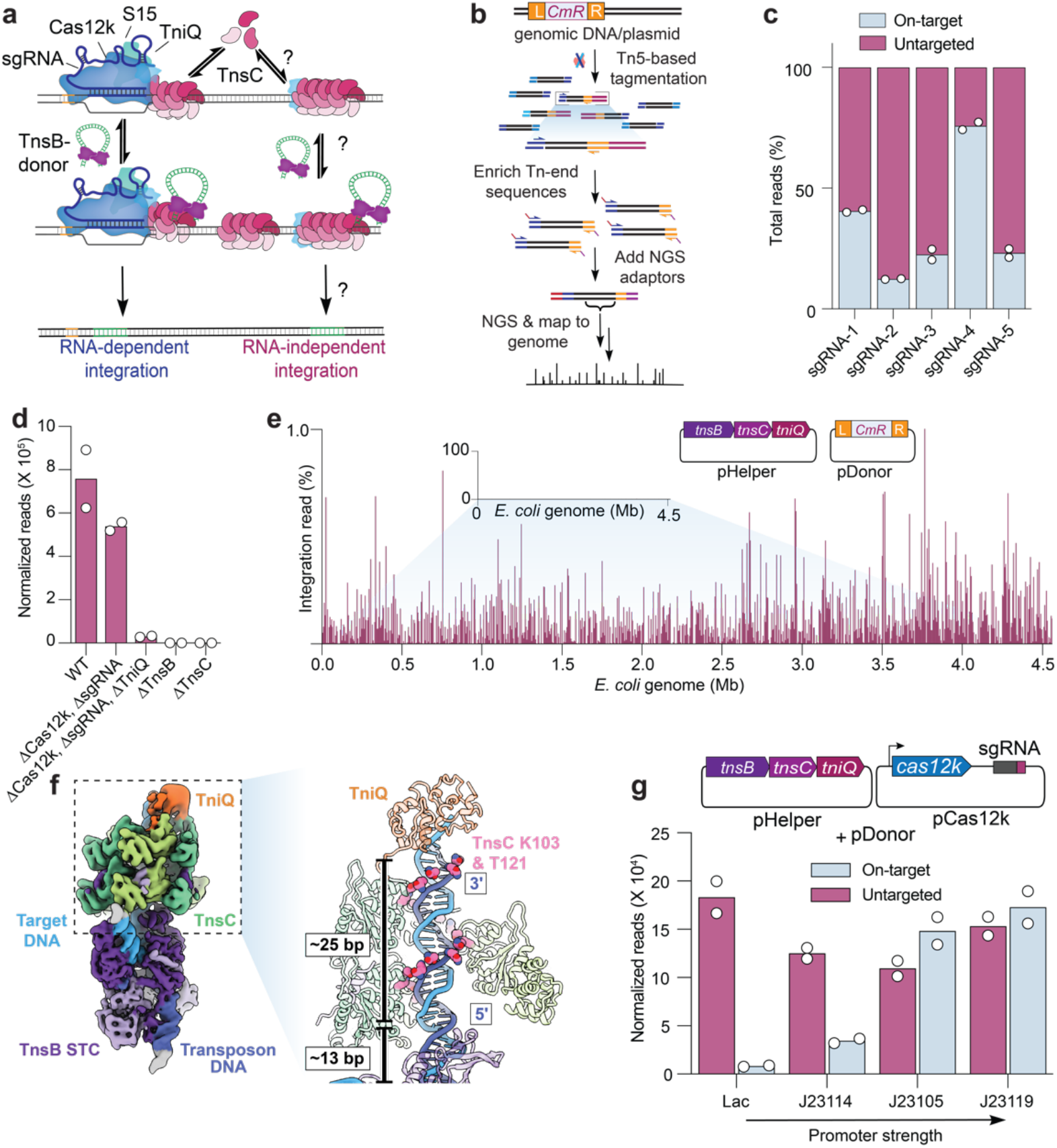
Type V-K CASTs direct frequent Cas12k- and RNA-independent transposition events. **a**, Schematic of type V-K CAST transposition occurring at on-target sites (RNA-dependent) and untargeted sites (RNA-independent). **b,** Experimental pipeline used for tagmentation-based transposon insertion sequencing (TagTn-seq) for *in vitro* and genomic samples. **c**, Fraction of total genome-mapping integration reads detected at on-target and untargeted sites for the wildtype pHelper expression plasmid across multiple guides (sgRNA-1 to sgRNA-5). **d,** Total genome-mapping reads detected for WT pHelper or pHelper with the indicated deletions, normalized and scaled (Methods). **e**, Zoomed-in view of integration reads comprising 1% or less of *E. coli* genome-mapping reads, in an experiment performed without the Cas12k and guide RNA. **f,** Cryo-EM reconstruction of the untargeted transpososome reveals the assembly of TniQ (orange), TnsC (green), and TnsB (purple) in a strand-transfer complex (STC). The target DNA and transposon DNA are represented in light blue and dark blue, respectively. For visualization, a composite map was generated using two local-resolution filtered reconstructions from the focused refinements (Methods). Zoomed-in and cutaway view showing TnsC forming a helical assembly on the target DNA, positioning residues K103 and T121 (pink) adjacent to one strand of the target DNA (dark blue). 5’ and 3’ ends of the TnsC-interacting DNA strand are indicated. Two turns of TnsC and TnsB footprint on DNA until TSD cover approximately 25 and 13 base pairs (bp), respectively. Only selected TnsC monomers are represented in the cutaway for clarity. **g**, Cas12k and the sgRNA were cloned onto a separate vector, and the promoter driving Cas12k expression was varied. Reads detected at on-target and untargeted sites during transposition assays were normalized and scaled (Methods). For **c**, **d, e**, and **g**, the mean is shown from n = 2 independent biological replicates.

Here we set out to rigorously investigate the mechanism of target site selection for the archetypal type V-K CAST system from *Scytonema hofmannii*, focusing special attention on the role of TnsC in regulating fidelity. Remarkably, we found that ShCAST is prone to extensive, RNA-independent transposition through a pathway that requires only TnsB, TnsC, and TniQ. Although these untargeted integration events initially appear random, analysis of high-throughput sequencing data revealed a significant bias for AT-rich sites, which was corroborated by single-molecule biophysical studies of TnsC DNA-binding behavior. By modulating DNA substrates in biochemical transposition assays, we demonstrated that the preference for AT-rich sequences could lead to predictable reaction outcomes. Furthermore, we found that transposition specificity could be substantially improved by limiting the cytoplasmic TnsC levels, further highlighting the role of TnsC filament formation in the pathway choice between RNA-dependent and RNA-independent transposition. Collectively, our results underscore the value of mechanistic studies in revealing new opportunities to engineer and leverage CAST systems as a potent DNA insertion technology.

## RESULTS

### Type V-K CASTs undergo RNA-dependent and RNA-independent transposition pathways

Previous studies of the type V-K ShCAST system from *Scytonema hofmannii* revealed that a considerable proportion of genomic integration events occur at sites distal from the target site dictated by the guide RNA^11, 14, 15^. To understand the molecular basis of these events, we applied a high-throughput sequencing approach to unbiasedly capture genome-wide integration events upon ShCAST expression with various genetic perturbations (**Fig. 1b, Methods**). After testing five distinct single-guide RNAs (sgRNA), we found that the fraction of on-target integration events ranged from 12–76%, and that the majority of events occurred elsewhere, with DNA insertions seemingly randomly distributed across the genome at low individual frequencies (**Fig. 1c**, **Supplementary Fig. 1a,b**). We analyzed their proximal genetic neighborhood and failed to detect enriched sequence similarity to the guide RNA (**Supplementary Fig. 1c-e**), suggesting that these events were not mismatched off-targets aberrantly targeted by RNA-guided Cas12k, but rather, the consequence of RNA-independent transposition; we therefore tentatively referred to these as untargeted integration events (**Fig. 1c**). When we deleted *cas12k* and the sgRNA from the original pHelper expression plasmid, the CRISPR-lacking ShCAST system still produced efficient genome-wide transposition products (**Fig. 1d,e**)^11^. These results clearly establish that type V-K CAST systems are capable of both RNA-dependent targeted DNA integration, and RNA-independent untargeted DNA integration.

We performed additional control experiments and confirmed that TnsC, a AAA+ regulator, and TnsB, the DDE-family transposase, are essential for both RNA-dependent and RNA-independent transposition, as their deletion completely abrogated integration (**Fig. 1d**). We initially hypothesized that TnsB and TnsC would comprise the minimum necessary protein components for RNA-independent transposition, similar to the reliance of phage Mu transposition on two homologous gene products, MuA and MuB^27^. To our surprise, *tniQ* deletion had a severe effect on untargeted transposition (**Fig. 1d**), suggesting a crucial role in stabilizing and/or interacting with the TnsBC transpososome. Recent structures reveal that DNA-bound TnsC oligomers are capped on the N-terminal face by one or more TniQ protomers^22, 26^, and our biochemical experiments similarly demonstrated that TniQ only stably associated with DNA in the presence of TnsC, as reflected by fluorescence polarization experiments (**Supplementary Fig. 1f**). Thus, much like the requirement for TnsB, TnsC, and TniQ in transposition by Tn*5053*^28^, we conclude that ShCAST — and perhaps type V-K CAST systems more generally — maintain a prominent ‘BCQ’ pathway that facilitates CRISPR RNA-independent, untargeted transposition.

To capture the architecture of components contributing to untargeted integration, we used cryo-electron microscopy (cryo-EM) to visualize TnsB, TnsC and TniQ in a strand-transfer complex (STC). Although the cryo-EM density of TniQ was less well-resolved (∼8 Å) compared to other subunits (**Supplementary Fig. 2**), likely due to heterogeneity of binding configurations, we were able to unambiguously dock atomic models of all protein components and DNA into the map (**Fig. 1f**). The overall structure of the ‘BCQ’ transpososome resembled the Cas12k-containing transpososome, with DNA-interacting residues of TnsC (K103 and T121) positioned to follow the helical symmetry of duplex DNA, as in the structure of helical TnsC^22–24, 26^. However, in contrast to the Cas12k-containing transpososome^23^, the polarity of the interacting DNA strand in the ‘BCQ’ transpososome was 3’ to 5’ following the direction of TniQ-to TnsB-binding face of TnsC, which was also noted in the structure of TniQ-TnsC^22, 26^. Therefore, the ‘BCQ’ transpososome structure, which represents a low energy configuration of TnsC, reveals that DNA contacts in TnsC filaments are maintained differently at on-target and untargeted sites. Yet, it is impressive that the overall architecture of the ‘BCQ’ transpososome, comprising the TnsB STC, two turns of a TnsC mini-filament (spanning a DNA-binding footprint of 25 bp) and TniQ, resembles the on-target Cas12k bound transpososome.

**Fig. 2.**
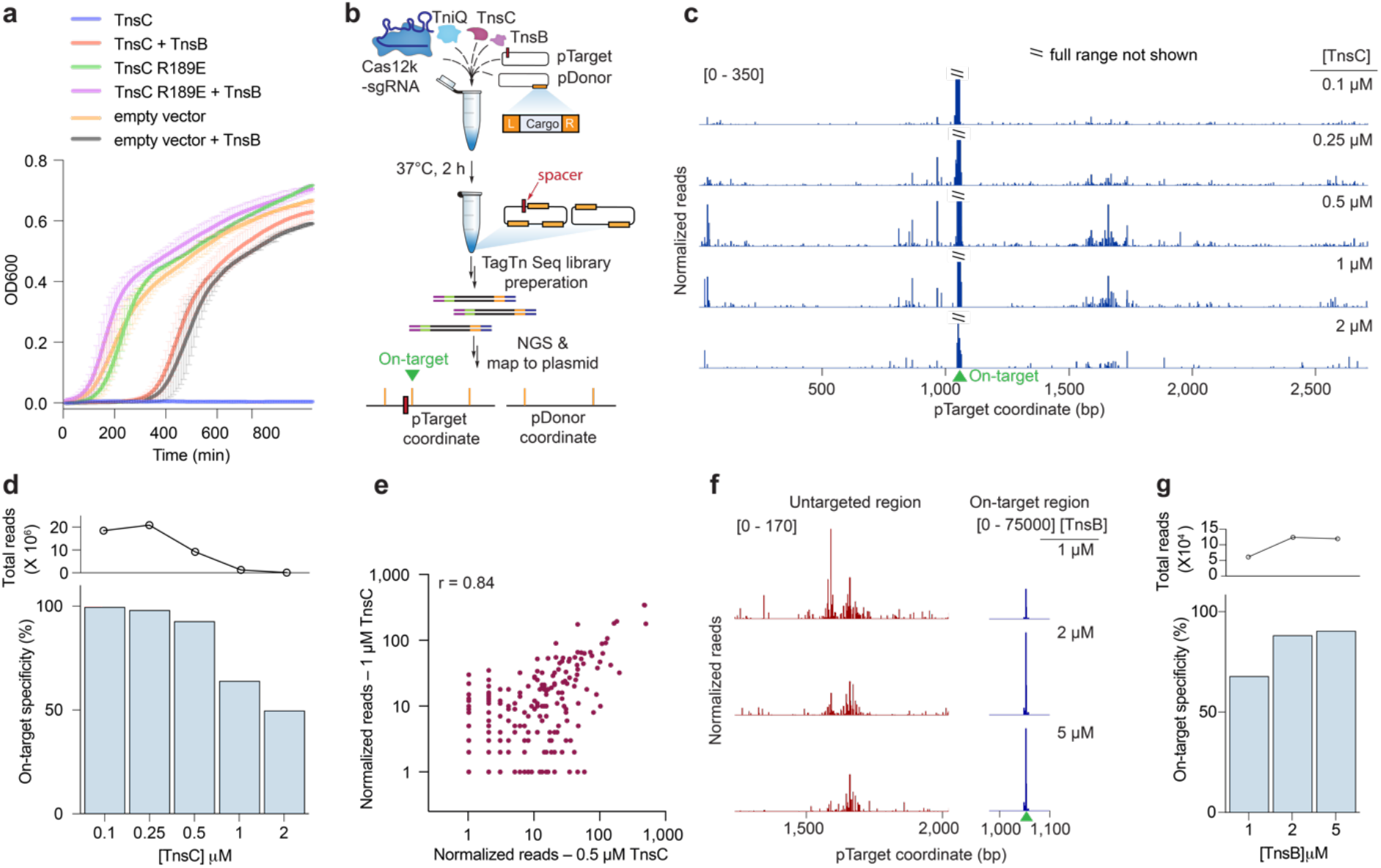
Biochemical reconstitution of transposition reveals distinct efficiencies at on-target and untargeted sites. **a**, Growth curves upon induction of WT or mutant TnsC, with or without TnsB. The data shown are mean ± s.d for n = 2 independent biological replicates, inoculated from individual colonies. **b**, Assay schematic for probing *in vitro* plasmid-to-plasmid transposition events using recombinantly expressed CAST components. **c**, In vitro integration reads mapping to pTarget, from experiments in which TnsC was titrated from 0.1 – 2 µM. Data were normalized and scaled to highlight untargeted integration events, relative to on-target insertions (**Methods**). **d**, On-target specificity from biochemical transposition assays at varying TnsC concentrations, calculated as the fraction of on-target reads divided by total plasmid-mapping reads (bottom). Total integration activity also decreased as a function of TnsC concentration, as seen by the normalized plasmid-mapping reads (top). **e,** Scatter plot showing reproducibility between untargeted integration reads observed *in vitro* at two high TnsC concentrations; each data point represents transposition events mapping to a single-bp position within pTarget. The Pearson linear correlation coefficient is shown (two tailed *P* <0.0001); on-target events were masked (**Methods**). **f**, Normalized integration reads detected at a representative untargeted site (left) and at the on-target site (right), with 1 µM TnsC and the indicated TnsB concentration. Note the differing y-axis ranges. **g**, On-target specificity from biochemical transposition assays at 1 µM TnsC and the indicated TnsB concentration, shown as in **d**.

We next wondered whether the presence of Cas12k and an appropriate sgRNA would reduce the frequency of untargeted transposition events by sequestering protein components at the on-target site. After cloning *cas12k* onto a separate expression plasmid and systematically varying its promoter strength, we found that on-target integration events were proportionally increased, though without a significant reduction in the frequency of untargeted integration (**Fig. 1g**, **Supplementary Fig. 1g**). These results indicate that, at least under these expression conditions, the availability of Cas12k-sgRNA complexes limits RNA-guided DNA integration efficiency but does not directly affect the ‘BCQ’ pathway. Interestingly, type V-K CAST systems often encode a MerR-family transcriptional regulator adjacent to the Cas12k gene^12, 29^, and a recent study demonstrated that these Cas V-K repressor (CvkR) proteins down-regulate both Cas12k and TnsB expression, though with distinct effects^30^. Thus, although our present knowledge about CAST activity is largely limited to comparative genomics and artificial heterologous over-expression, it appears likely that CAST transposition in native contexts is regulated so as to modulate the frequency of RNA-dependent and RNA-independent target pathways.

### Relative TnsB and TnsC stoichiometry determines the transposition pathway choice

Many other bacterial transposons encode transposition proteins homologous to ShCAST, including type I CASTs, Tn*7*, Tn*5053*, IS*21*, and Mu^4, 10, 27, 28, 31^. The TnsBC module is common to all, and in the case of Mu, the AAA+ ATPase component, known as MuB, plays a dominant role in directing untargeted, genome-wide transposition by recruiting the MuA transposase to potential integration sites^5^. Moreover, structural studies have demonstrated that MuB and ShTnsC both form continuous, non-specific filaments on double-stranded DNA (dsDNA)^5, 24, 26^, in contrast to the discrete, closed or semi-closed rings formed by TnsC from VchCAST (Tn6677) and *E. coli* Tn*7*^32, 33^. Therefore, we set out to experimentally investigate the role of TnsC in target site selection and the effect of variable stoichiometries of TnsB, TnsC and TniQ on untargeted integration. However, one of the major hindrances we encountered while trying to vary the expression of transposon components in cells was the toxicity associated, particularly with the over-expression of TnsC (**Fig. 2a**, **Supplementary Fig. 3a,b**). When we inoculated liquid cultures with a strain expressing *tnsC* alone from a strong promoter and induced over-expression in the lag phase, we observed a complete growth arrest for the majority of clones, with only a few strains undergoing delayed growth, likely due to suppressor mutations in the plasmid or genome (**Supplementary Fig. 3a)**. Strikingly, this cellular toxicity was completely rescued with a mutation to the arginine finger motif, which abrogates TnsC filamentation^34^ and transposition, or partially rescued by co-expression of TnsC and TnsB (**Fig. 2a**, **Supplementary Fig. 3c**). These results implicate non-specific DNA filamentation as a likely source of cellular toxicity, which can be relieved in part by the ability of TnsB to disassemble TnsC filaments, as demonstrated from in vitro experiments^24, 26, 35^.

**Fig. 3.**
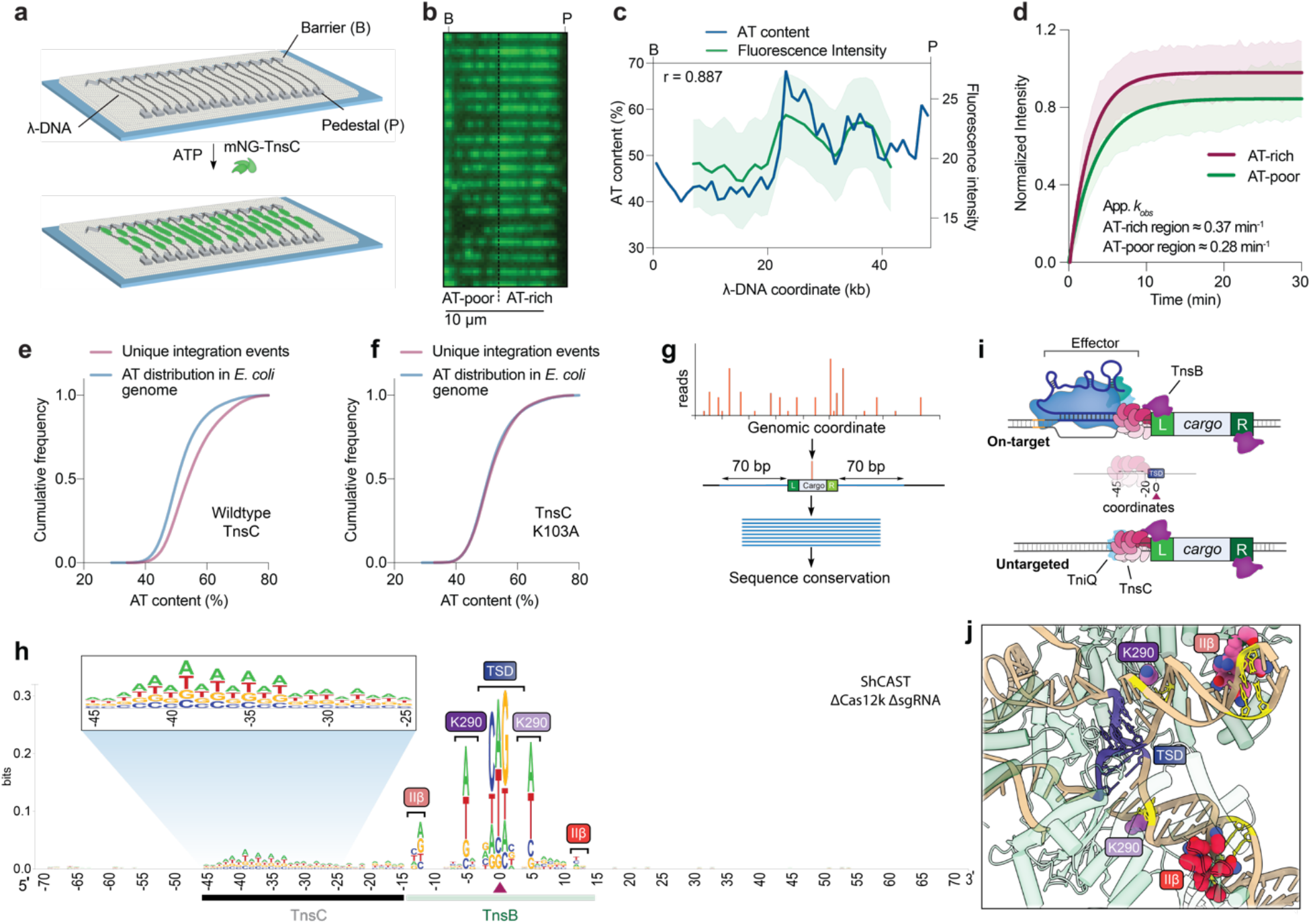
RNA-independent integration events occur at preferred sequence motifs. **a**, Schematic for single-molecule DNA curtains assay to visualize TnsC binding. λ phage DNA substrates are double-tethered between chrome pedestals and visualized used TIRF microscopy. **b**, mNG-labeled TnsC preferentially binds AT-rich sequences on the λ-DNA substrate near the 3’ (pedestal) end (**Supplementary Movie 1**). **c**, Correlation between AT content and mNG-TnsC fluorescence intensity visualized along the length of λ DNA. The Pearson linear correlation coefficient is shown (two-tailed *P* < 0.0001); data shown represent the mean ± s.d for n = 66 molecules. **d**, Binding kinetics for mNG-TnsC at AT-rich and AT-poor regions of the λ-DNA substrate. Apparent *k_obs_* at AT-rich sites ≈ 0.37 min^-1^, 95% C.I. [0.35, 0.39] and at AT-poor sites ≈ 0.28 min^-1^, 95% C.I. [0.27, 0.30]. The data shown represent mean ± s.d for n = 87 molecules. **e,** Cumulative frequency distributions for the AT content within a 100-bp window flanking integration events, using ShCAST with WT TnsC and sgRNA-1 (n = 5,505 unique integration events), compared to random sampling of the *E. coli* genome (n = 50,000 counts; **Methods**). The distributions were significantly different, based on results of a Mann-Whitny *U* test (*P* = 1.48 x 10^-135^). **f,** Cumulative frequency distribution comparison as in panel **e**, but with a K103A TnsC mutant (n = 1,932 unique integration events), which revealed a loss of AT bias (*P* = 0.1349). **g,** Meta-analysis of untargeted transposition specificity was performed by extracting sequences from a 140-bp window flanking the integration site and generating a consensus logo. **h,** WebLogo^60^ from a meta-analysis of untargeted genomic transposition (n = 5,855 unique integration events) with a modified pHelper lacking Cas12k and sgRNA. The site of integration is noted with a maroon triangle. An AT-rich sequence spanning ∼25 bp likely reflects the footprint of two turns of a TnsC filament (black), whereas motifs within/near the target-site duplication (TSD) represent TnsB-specific sequence motifs (green). Specific TnsB residues/domains contacting the indicated nucleotides are shown. The zoomed-in inset highlights periodicity in the sequence bound by TnsC. **i,** Schematic showing the relative spacing of sequence features bound by Cas12k, TnsC, and TnsB in both on-target (RNA-dependent) and untargeted (RNA-independent) DNA transposition. In both cases, the TnsC footprint covers ∼25-bp of DNA and directs polarized, unidirectional integration downstream in a L-R orientation. **j**, Zoom-in view of the ShCAST transpososome structure, highlighting sequence-specific contacts between TnsB and the target DNA that were observed in the WebLogo in **h**. PDB ID: 8EA3^23^.

In order to modulate the stoichiometries of CAST components contributing to untargeted integration, while avoiding confounding factors such as toxicity, we adopted a biochemical approach. After recombinantly expressing and purifying ShCAST components and testing the activity of TnsC and TnsB *in vitro* (**Supplementary Fig. 3d-g**), we established a plasmid-to-plasmid (pDonor-to-pTarget) transposition assay (**Fig. 2b**, **Supplementary Fig. 3d-g**). In initial experiments, we amplified on-target products by targeted PCR, thereby revealing molecular requirements for each of the transpososome components and the expected distance separating the target and integration site (**Supplementary Fig. 4a-c**). Next, we coupled our biochemical experiments with tagmentation-based high-throughput sequencing, in order to unbiasedly map DNA transposition events regardless of their insertion site (**Fig. 2b, Methods**). Remarkably, at low (0.1 µM) concentrations of TnsC, transposition was highly accurate, with >99% of reads representing on-target integration events, defined as occurring within a 100-bp window downstream of the target site (**Fig. 2c,d**). However, when we systematically increased the concentration of TnsC while keeping all other components constant, the frequency of untargeted integration events increased as well, approaching levels similar to those observed in cellular transposition assays (**Fig. 2d**, **Supplementary Fig. 4e**). Substantial untargeted integration events also occurred in the absence of Cas12k and sgRNA under these conditions, in agreement with *in vivo* experiments (**Supplementary Fig. 4d**). When interpreted together with structural data, these results suggest that RNA-independent transposition is likely initiated by the formation of dsDNA-bound TnsC filaments. Importantly, untargeted integration events were not randomly distributed across pTarget but instead clustered into specific and reproducible hotspot regions (**Fig. 2e**), suggesting a selectivity for certain, yet undetermined sequence features (see below).

**Fig. 4.**
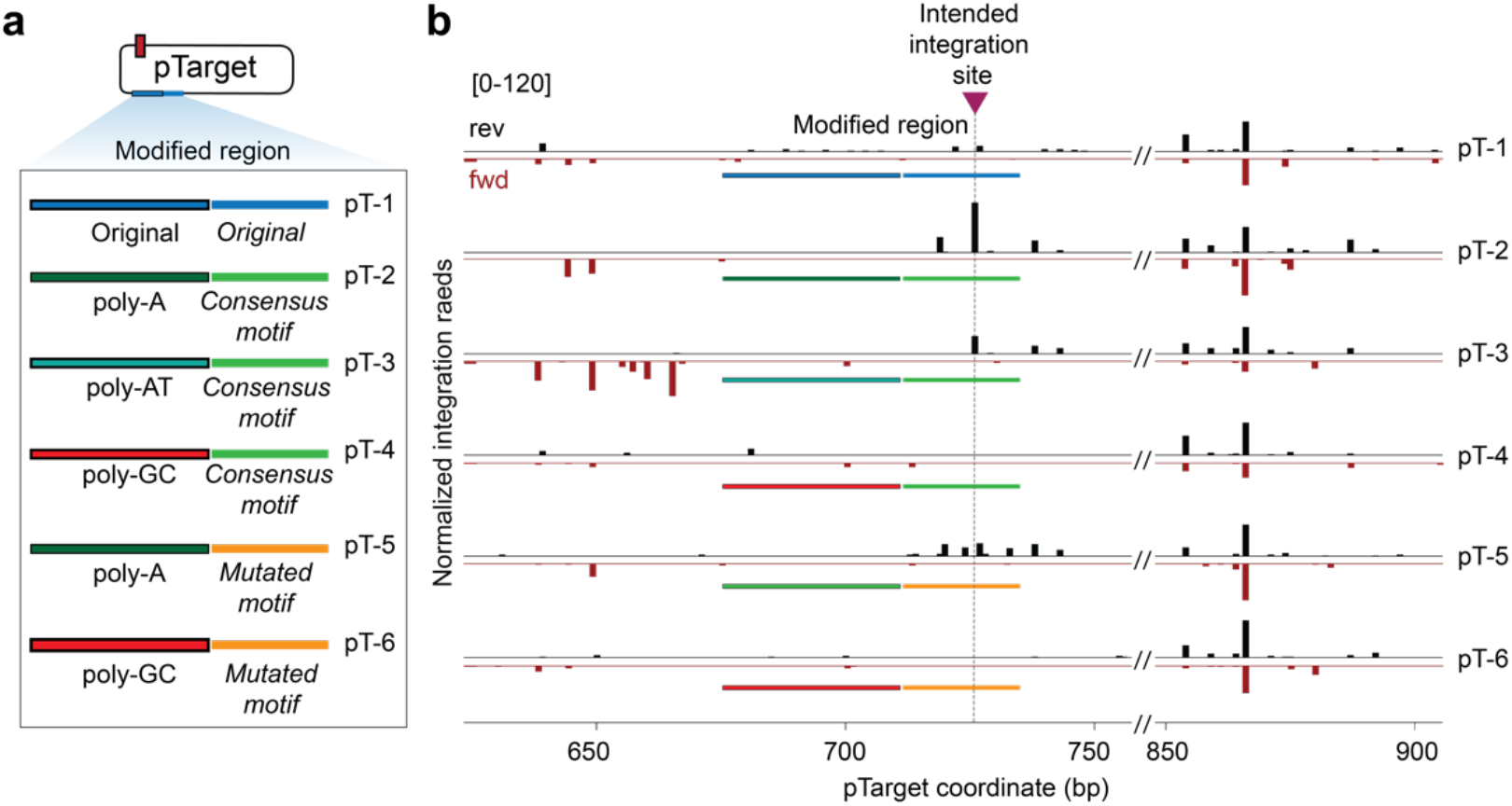
Artificial induction of semi-targeted RNA-independent transposition at preferred motifs. **a**, A region on pTarget exhibiting low integration activity (original, blue) was substituted with rationally engineered sequences (colored) based on TnsC and TnsB binding preferences, generating the indicated pTarget variants (pT-1–6). **b,** After performing biochemical transposition assays with the indicated pTarget substrates, integration reads were normalized and mapped to either the forward strand (fwd, red) or reverse strand (rev, black). The intended ‘untargeted’ integration site based on optimized poly-A and TnsB consensus motifs is marked with a maroon triangle and dotted line; the representative region at right (850–900 bp) is shown to highlight consistency in integration events observed elsewhere on pTarget.

We next tested the impact of other protein components on *in vitro* transposition activity. Ribosomal protein S15, a recently described host factor that stimulates ShCAST transposition by binding the sgRNA^22^, dramatically increased the frequency of on-target integration events, as measured both by deep sequencing and qPCR, but had no discernible effect on untargeted integration events (**Supplementary Fig. 4g,h**). Increasing the concentration of TniQ, on the other hand, led to a monotonic increase in the frequency of untargeted integration events without a major effect on on-target integration (**Supplementary Fig. 4f**), suggesting that the RNA-independent pathway may be more sensitive to limited TniQ availability.

The TnsB transposase has been previously shown to disassemble TnsC filaments from dsDNA^24, 26, 35^, and it is therefore possible that titrating excess amounts of TnsB would lead to partial or full disassembly of TnsC filaments necessary for transposition, regardless of their molecular context. Yet, when we varied the amount of the TnsB transposase, we observed distinct effects at on-target and untargeted sites (**Fig. 2f**, **Supplementary Fig. 4i-k**). Increasing TnsB led to a notable increase in RNA-guided integration but resulted in a slight decrease in untargeted events (**Supplementary Fig. 4i-k)**, leading to an overall rescue of specificity at on-target sites with high TnsC concentrations. This observation suggests that TnsC filaments at targeted versus untargeted sites are differentially susceptible to TnsB-induced disassembly, and/or react to undergo strand transfer with distinct kinetics. Transpososome structures reveal that TnsB interacts with TnsC filaments on only one face^23, 35^, and no major structural changes are associated with TnsC filaments at on-target and untargeted sites (**Fig. 1f**). Therefore, we suggest that the distinct nature of DNA interactions made by TnsC at both these sites determine filament stabilization versus disassembly.

Collectively, these results suggest that the natural propensity of TnsC to form long filaments on dsDNA exerts a fitness cost on cells in the absence of accessory transposase machinery and is a driver of RNA-independent, untargeted transposition. We next sought to investigate whether TnsC exhibits any bias when selecting RNA-independent sites for transposition.

### TnsC preferentially targets AT-rich DNA during RNA-independent transposition

We developed a single-molecule approach to visualize DNA binding by fluorescently-labeled TnsC using DNA curtains (**Fig. 3a**, **Supplementary Fig. 5a, Supplementary Movie 1**), in which λ-phage genomic DNA molecules are tethered between chrome patterns on a quartz slide and imaged by total internal reflection microscopy (TIRFM)^36^. TnsC exhibited stable and high-affinity binding in the presence of ATP, and the data furthermore revealed a marked preference for the 3’ half of the genome (**Fig. 3b**). The λ-phage genome is known to be divided into a GC-rich half and an AT-rich half^37^, and our analyses revealed a significant correlation between AT content and TnsC localization (**Fig. 3c**), indicating that TnsC filaments preferentially accumulated on the AT-rich half of the λ-phage genome. Time-course experiments further revealed that TnsC binds to AT-rich regions at a faster rate, before saturating the entire λ-DNA substrate within 5-10 min of incubation (**Fig. 3d**, **Supplementary Fig. 5a,b**). Interestingly, a preference for AT-rich regions has been previously observed for MuB, with both single-molecule microscopy experiments and *in vivo* transposition studies^38, 39^, supporting the idea that this property is likely to be broadly conserved across AAA+ regulators from other transposon families.

**Fig. 5.**
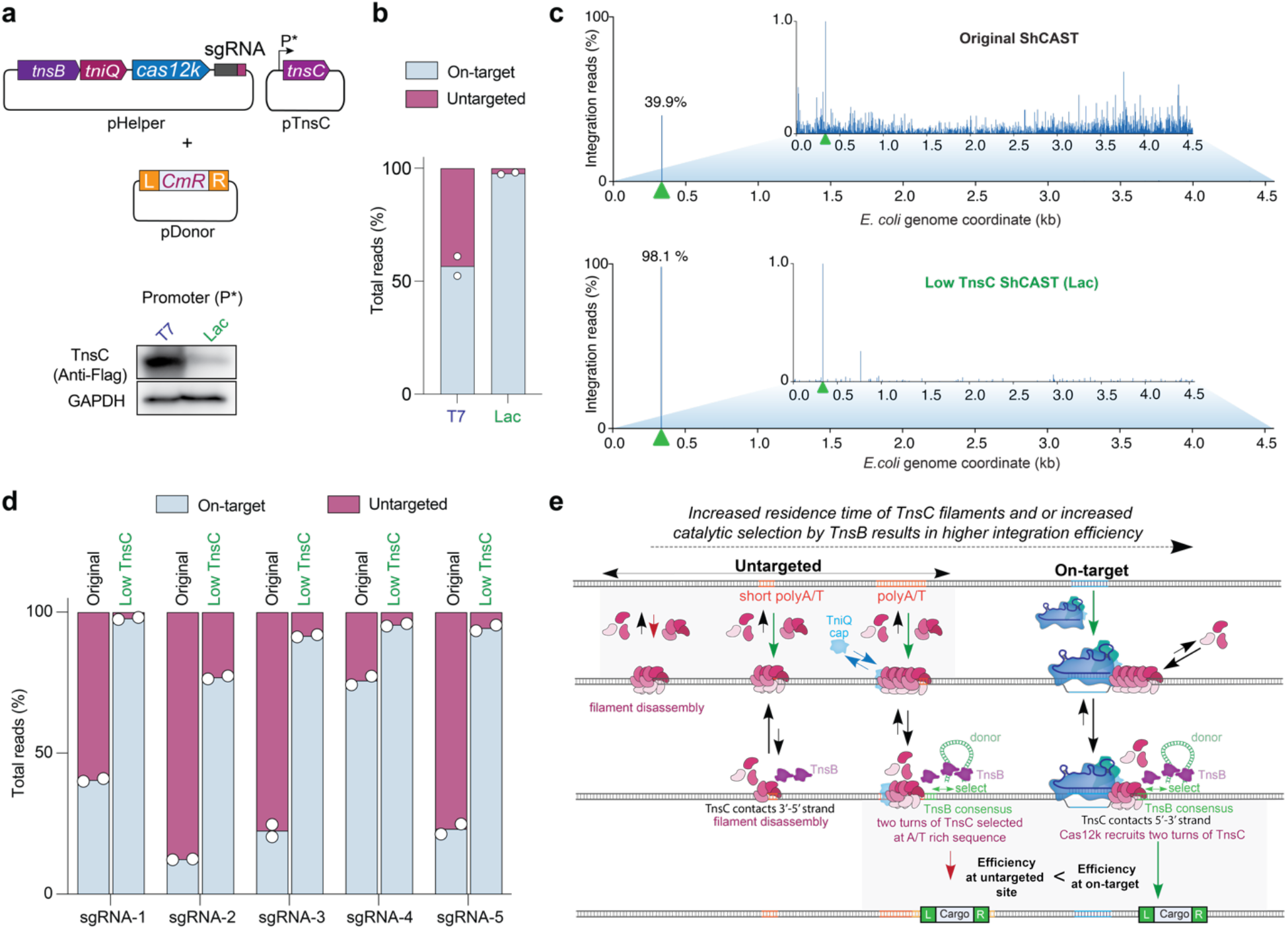
The fidelity of RNA-guided DNA integration is controlled by TnsC concentration. **a**, Schematic of alternative ShCAST expression strategy, in which TnsC was encoded on a separate plasmid (pTnsC) driven by a Lac or T7 promoter. Distinct cellular expression levels were confirmed by Western blot against a 3xFLAG epitope tag fused to TnsC (bottom). **b**, Fraction of total genome-mapping integration reads detected at on-target and untargeted sites upon TnsC expression with a Lac or T7 promoter. **c**, Genome-wide view of *E. coli* genome-mapping reads for the original/WT ShCAST system as compared to a modified ShCAST system with low TnsC expression; the zoomed-in view visualizes reads comprising 1% or less of genome-mapping reads. The target site is marked with a green triangle. **d,** Fraction of total genome-mapping integration reads detected at on-target and untargeted sites, with the original ShCAST system or modified ShCAST system with low TnsC expression. Data for five sgRNAs are shown. For **b** and **d**, the mean is shown from n = 2 independent biological replicates. **e,** Model for target-site selection and transpososome assembly during on-target, RNA-dependent transposition (right) or untargeted, RNA-independent transposition (left) by type V-K CAST systems. Within the untargeted pathway, TnsC preferentially forms filaments at A/T-rich regions and is capped by TniQ, leading to the downstream site being selected by TnsB for integration. Cas12k-bound targets may better nucleate TnsC filament formation, and we hypothesize that TnsC filaments loaded at Cas12k-bound targets serve as better substrates for DNA integration, compared to untargeted sites. Importantly, all structures of TnsC filaments representing untargeted sites^22–24, 26^ including the ‘BCQ’ transpososome, reveals K103 residues of the TnsC monomers forming the filament proximal to TnsB, contacting DNA with opposite strand polarity compared to on-target structures^22, 23^. This could be decisive for the distinct efficiencies observed at these sites.

Given our observation that untargeted transposition events in biochemical assays preferred certain hotspot regions of pTarget and were reproducible between independent experiments (**Fig. 2e**), we hypothesized that AT content might be an underlying feature explaining these data. We analyzed the nucleotide composition surrounding all unique integration events on pTarget and found that they were indeed skewed towards more AT-rich DNA (**Supplementary Fig. 5d,e, Methods**). Direct visual superposition of AT content and DNA integration data further revealed that hotspot regions for untargeted transposition in pTarget generally correlated with regions of higher AT content (**Supplementary Fig. 5f)**. We observed the same phenomenon after performing transposition assays with a λ-DNA substrate and repeating similar analyses to assess AT-bias in the location of untargeted integration sites (**Supplementary Fig. 5g-i**). Finally, we analyzed untargeted integration events in the *E. coli* genome, from experiments performed without Cas12k and sgRNA, and found that these were also highly enriched at local regions of high AT content (**Fig. 3e**, **Supplementary Fig. 5j**). These results provide strong evidence that AT-rich sites on DNA are preferentially bound by TnsC, and thus preferentially ‘targeted’ for RNA-independent transposition.

We next sought to uncover additional sequence features common to CRISPR-independent ShCAST transposition products. We performed a meta-analysis of all genome-wide integration sites after orienting the flanking sequences based on the asymmetric transposon ends, and then generated a consensus sequence logo of the resulting alignment (**Fig. 3g, Methods**). This analysis revealed two notable clusters of sequence features: nucleotide preferences directly within and surrounding the target-site duplication (TSD), and an AT-rich nucleotide cluster located upstream of the integration site (**Fig. 3h**). Remarkably, the AT-rich region spans ∼25-bp and could thus accommodate two turns of a dsDNA-bound TnsC filament, similar to the TnsC architecture and footprint observed within the context of Cas12k-containing and Cas12k-lacking transpososomes (**Fig. 1f,g**)^23^. The observation that this region is located on only one side of all integration sites strongly argues that RNA-independent integration events result from binding of TnsC filaments to AT-rich DNA, followed by directional recruitment of TnsB to define downstream sites for transposon insertion in the same L-R orientation as occurs at RNA-guided target sites^11^ (**Fig. 3i**). In general, we refer to this orientation as TnsC-LR which could be applicable to other systems inserting employing a AAA+ ATPase. Interestingly, the highest sequence conservation was located furthest from the site of integration, proximal to the presumed region where TnsC filaments are capped by TniQ (**Fig. 1f,g**), and the observed, di-nucleotide periodic trend is reminiscent of structures demonstrating that TnsC monomers contact every two bases of DNA^24, 26^.

The highest conservation in the sequence logo corresponds to sequences contacted by TnsB within the ‘BCQ’ transpososome. As with prior library-based experiments for both type I-F and V-K CAST systems, our results indicate that TnsB preferentially integrates into sites containing ‘GCWGC’ within the TSD (**Fig. 3h**)^11, 40^. However, we also uncovered a novel bias for (A/T) at symmetric positions located ± 5-bp from the TSD center, which is contacted by residue K290 of two TnsB monomers within the transpososome (**Fig. 3j**). Nucleotide preferences ± 12-bp from the TSD could also be explained by the proximity of these residues with the TnsB IIß DNA-binding domain (R416, T417, Q425, N428), which importantly, also makes similar sequence contacts to the penultimate TnsB binding sites located within the transposon left and right ends^23, 35, 41^. When we analyzed untargeted integration events from our previously published ShCAST data^14^, in which HTS libraries were generated and sequenced using an alternative strategy, we observed the exact same sequence features, confirming the robustness of this observation (**Supplementary Fig. 6a**). Importantly, the absence of any conserved sequence features upstream of the A/T-rich region, where the target site would normally be located during Cas12k-mediated transposition, corroborated our earlier interpretation that the majority of cellular transposition events are RNA-independent.

Previously, it was shown that a point mutation in one of the two TnsC (K103A) residues that contact DNA increased the number of untargeted events without comprising its ability to bind DNA^26^. In agreement with these results, when we tested the same mutant in cellular transposition assays, we observed a severe loss of on-target events but a preservation of untargeted events, which were enriched near the *E. coli* origin of replication (**Supplementary Fig. 3c, 5k**). Strikingly, when we performed a meta-analysis of untargeted integration events, we found that the TnsC K103A mutant no longer exhibited an A/T preference, in contrast to WT TnsC (**Fig. 3f**, **Supplementary Fig. 6b**). This observation, together with the loss of on-target integration (**Supplementary Fig. 3c**), suggests that the K103A mutation results in a more promiscuous mode of DNA binding that supports integration anywhere in the genome without specific sequence requirements.

We previously reported transposition activity for a type V-K CAST homolog also found in *Scytonema hofmannii*, ShoCAST (previously referred to as ShoINT), which is diverged from ShCAST and more similar to AcCAST^11, 14^. We were curious as to whether untargeted ShoCAST transposition events would exhibit similar sequence preferences as ShCAST and thus performed meta-analyses on published transposition data^14^. Highly similar motifs emerged in the resulting sequence logo, but with a major difference in the window of A/T-rich DNA located upstream of the integration site, which spanned only ∼10-bp compared to ∼25-bp observed with ShCAST (**Supplementary Fig. 6c**). This difference is remarkably consistent with the finding that ShoCAST and AcCAST integrate ∼10-bp closer to the target site than ShCAST^11, 14^, suggesting that the transpososomes from this subfamily of CAST systems — for both RNA-dependent and RNA-independent transposition pathways — may comprise a shorter TnsC filament spanning only one turn of DNA.

Altogether, these observations demonstrate how subtle sequence motifs at RNA-independent integration sites can be gleaned from meta-analyses of genome-wide integration data. They furthermore clearly reveal that RNA-independent transposition is not random, but rather, that the ‘BCQ’ transposition pathway preferentially selects certain genomic regions over others.

### Preferred sequence motifs lead to semi-targeted, RNA-independent transposition

To test the importance of TnsBC-specific sequence motifs more directly for CRISPR-independent integration, we designed biochemical transposition assays using a series of isogenic pTarget substrates that differed only in the sequence content of a select region that was poorly targeted in previous experiments (**Fig. 4a**, substrate pT-1). We hypothesized that we could generate ‘targeted’, RNA-independent insertions within this region if an optimal sequence were designed to include both the poly-A stretch and flanking TnsB consensus motifs observed in the sequence logo described above (substrate pT-2; **Fig. 3h**). As further controls, we substituted the poly-A stretch with either poly-AT or poly-GC, we mutagenized the TnsB consensus, or we replaced both motifs (**Fig. 4a**, substrates pT-3 through pT-6). We then tested each substrate in biochemical transposition assays and plotted the normalized integration frequency within this window of interest.

The resulting data demonstrate that RNA-independent integration events occur in predictable ways depending on the sequence features uncovered through our analyses (**Fig. 4b**). Substrate pT-2 exhibited a predominant integration product precisely at the engineered site, in the expected T-LR orientation, whereas this integration product was entirely absent when the poly-A was replaced with poly-GC, strengthening our conclusion that favorable TnsC filamentation is important for RNA-independent integration (**Fig. 4b**, pT-4,6). When we retained the poly-A stretch but mutated the consensus motif favored by TnsB, integration products were more heterogeneously positioned (**Fig. 4b**, pT-5), suggesting that preferred TnsB-DNA interactions play an important role in dictating the exact insertion site, as similarly concluded by our recent study on the type I-F VchCAST system^40^. Interestingly, when we replaced the poly-A sequence with poly-AT (increasing the A-content in the opposite strand), the intended integration event was diminished in frequency and accompanied by an increase in upstream integration events on the opposite strand (**Fig. 4b**, pT-3), demonstrating that nucleotide composition can modulate the preferred directionality of TnsC filament formation and thus integration.

These experiments reveal that TnsC prefers to filament unidirectionally on A-rich DNA stretches, leading to downstream integration in the T-LR orientation. The efficiency and exact site of integration is thus a combination of TnsC filament formation propensity and local TnsB sequence preferences.

### TnsC availability controls the specificity of cellular ShCAST transposition activity

Beyond highlighting the role of TnsC in biasing untargeted integration events to occur at select hotspot regions of the genome, our results more generally implicate TnsC filament formation as a major driver of RNA-independent transposition activity. Because our *in vitro* results suggest that TnsB differentially selects TnsC filaments at on-target versus untargeted sites, we hypothesized that this difference could be exploited to increase the overall on-target integration accuracy. To test this hypothesis, we designed perturbations intended to repress TnsC filament formation at non-Cas12k-bound target sites, either by fusing TnsC directly to CRISPR effector proteins or by lowering overall TnsC expression levels.

When we fused Cas12k and TnsC, we observed an increase in on-target accuracy (**Supplementary Fig. 7a**), as was recently reported by Tou *et al*^20^. We initially hypothesized that this effect might result from local seeding of TnsC filaments upon Cas12k target binding, but co-expression of unfused Cas12k had no adverse effect on specificity, suggesting instead that TnsC filamentation may be partially impaired with an N-terminal adduct. We also replaced Cas12k with dCas9 and generated dCas9-TnsC fusions, hoping to similarly seed TnsC filaments at target sites bound by dCas9-sgRNA complexes. However, we noted no observable on-target integration and severely diminished untargeted integration events (**Supplementary Fig. 7b**), suggesting these designs were non-functional. Altogether, these experiments suggested that fusion strategies may be poorly suited to increase the probability of TnsC filament formation at RNA-dependent target sites without extensive further engineering and mutagenesis.

Next, we pursued an alternative strategy, motivated by our biochemical observation that increasing TnsC concentration tilted the balance between RNA-dependent (on-target) and RNA-independent (untargeted) transposition towards the latter pathway **(Fig. 2c,d**). To test if the same feature was applicable in cellular experiments, we relocated *tnsC* from the original high-copy pHelper plasmid to a separate, medium-copy plasmid where it was controlled by its own promoter (**Fig. 5a**). When we tested genomic integration activity under various promoter strengths, we observed dramatic differences in on-target specificity (**Fig. 5a,b**). Consistent with our *in vitro* results, low TnsC expression from a lac promoter resulted in 98% of integration events occurring on-target, whereas high TnsC expression with a T7 promoter resulted in considerably lower accuracy (57%), akin to the original pHelper vector (**Fig. 5a-c**, **Supplementary Fig. 7c,d**). Interestingly, cells expressing TnsC under control of a T7 promoter also showed a significant enrichment for insertion events across the T7 RNAP gene, suggesting that these clones were likely enriched within the population as a way of escaping TnsC-induced toxicity (**Supplementary Fig. 7c,e)**.

To determine whether this increased specificity effect was generalizable, we tested a range of guides previously shown to exhibit low on-target accuracy when tested with pHelper and pDonor. In all cases, we observed a substantial increase in the relative frequency of on-target integration events with the modified CAST construct (**Fig. 5d**). Importantly, this effect did not come at the expense of efficiency, as qPCR measurements revealed that on-target integration occurred with an equal or higher efficiency under low TnsC conditions, as compared to our original pHelper design (**Supplementary Fig. 7f**). It is possible that at low TnsC conditions, the decreased availability of TnsC at untargeted sites presumably titrates fewer TnsB-DNA complexes away from on-target sites and might be causative to the increased on-target efficiency.

Collectively, these results highlight the importance of relative expression levels for distinct components when delivering CAST machineries into target cells of interest, and further confirm the key role of TnsC in driving RNA-independent transposition events.

## DISCUSSION

Recent structures have shed light on the assembly of transpososome components for RNA-guided integration by CAST systems^22, 23^. However, a major proportion of integration events for type V-K CASTs occur at untargeted sites across the genome, for which there was no known mechanistic basis. Combining structural and functional evidence, we establish that type V-K CASTs maintain a distinct RNA-independent pathway facilitated by TnsB, TnsC, and TniQ (**Fig. 5e**). Our experiments reveal that the ability of TnsC to promiscuously form filaments on AT-rich DNA is a major driver of untargeted insertions. The role of TnsB in transposition, particularly at untargeted sites, is somewhat paradoxical, given its ability to disassemble TnsC and simultaneously facilitate integration. We speculate that stochastic TniQ binding might stabilize a specific configuration of TnsC filaments at untargeted sites, making them resistant to TnsB-mediated dissociation and instead promoting strand-transfer. TnsC disassembly may therefore be less efficient in cellular contexts than originally observed *in vitro* at high protein concentrations^24, 26, 35^. Our results highlight the competition between TnsB recruitment at TnsC-bound RNA-guided target sites versus AT-rich untargeted sites and indicate that TnsB preferentially reacts with Cas12k-bound on-target sites compared to untargeted sites (**Fig. 5e**). Future work will be necessary to resolve more precise kinetics of TnsC filamentation/disassembly in the presence of TniQ and TnsB, and differential TnsB transposition kinetics as a function of TnsC assembly state.

The structure of the ‘BCQ’ strand-transfer complex reveals two turns of TnsC filaments, an overall architecture reminiscent of the Cas12k-containing on-target transpososome^23^, with no major structural differences associated with TnsC in either of these assemblies. However, TnsC residues K103 and T121 proximal to TnsB in the ‘BCQ’ transpososome contact the DNA in 3’ to 5’ strand polarity, following the direction of TniQ- to TnsB-binding face of TnsC (**Fig. 1f**). The same strand is contacted in the case of random TnsC-DNA filaments^24, 26^ and the ‘non-productive’ Cas12k transpososome^22^, whereas in the productive on-target Cas12k transpososome, TnsC monomers proximal to TnsB contact the opposite strand (5’ to 3’). This interaction is thought to be a consequence of TnsC filament nucleation by Cas12k-TniQ and stabilization by additional DNA interactions (R182 and K119)^22, 23^. TnsC variants with mutations to the DNA-contacting residues retain the ability to filament on DNA^24, 26^, and maintain a significant proportion of integration at untargeted sites accompanied by a drop in on-target integration (**Supplementary Fig. 3c**). This suggests that these residues may not be a prerequisite for TnsC-DNA binding. Rather, we propose that these residues may serve as an intrinsic regulatory feature to ensure that random TnsC filaments default to contacting in the 3’ to 5’ strand polarity. This interaction mode could represent an energetically less favored or ’passive’ TnsC configuration for TnsB-mediated integration, thereby permitting only a subset of sites scanned by TnsC in the genome to be licensed for untargeted transposition.

It is well known that poly-A tracts in the genome represent regions of altered DNA curvature^42^, and our single-molecule experiments reveal that TnsC filamentation exhibits inherent affinity for AT-rich locations. Further meta-analyses of integration data revealed a preference for AT-rich sequences across a ∼25-bp window spanning ∼2 turns of a TnsC filament, upstream of features recognized by TnsB. We suggest that AT-rich genomic regions with altered DNA curvature may resemble the bending of DNA observed between unproductive and productive Cas12k transpososomes^22^, leading to preferential TnsC recruitment and a more favorable DNA binding mode that promotes TnsB-based DNA integration. To our surprise, the TnsC K103A mutation led to a complete loss of AT preference in the integration profile, suggesting that this mutant may achieve an energetically more favorable filamentation state, regardless of nucleotide composition. The loss of on-target integration for K103A may result from mutant TnsC filaments titrating TnsB-donor DNA complexes to untargeted sites in the genome more effectively than WT TnsC filaments.

The type V-K ‘BCQ’ pathway, although not exactly similar, resembles the TnsE-mediated pathway in Tn*7*-like transposable elements, in which structural features associated with DNA replication are recognized to primarily drive widespread mobilization into plasmids^7, 8^. Importantly, a gain-of-function TnsC mutant (A225V) was also identified for *E. coli* Tn*7*, which was capable of transposition in the absence of either of the two targeting factors, TnsD and TnsE^43^, and which facilitated integration at AT-rich sequences^44^. It is possible that in a native setting, type V-K CASTs exhibit low-frequency insertion at AT-rich sites, and that might be triggered by certain stimuli specific to cyanobacteria. Such a model would imply a transient selfish behavior by CASTs, possibly when the availability of plasmid-targeting guide RNAs are limiting for its proliferation or in situations when mobilization to a new AT-rich site is beneficial for propagation of the element. Although there is currently no direct evidence to support this hypothesis, recent experiments with a native type V-K CAST system in cyanobacteria indicates that expression of Cas12k and TnsB are regulated by a CvkR transcriptional repressor^30^, and thus could be important in modulating the choice between targeted and untargeted transposition pathways. Considering the highly conserved operonic nature of TnsB, TnsC, and TniQ in Tn*7*-like elements, Tn*5053*, and type V-K CASTs, tight regulation in the stoichiometry of these proteins could be important for accessing an RNA-independent untargeted pathway^13^. Future studies will be necessary to investigate this hypothesis further, by deep sequencing bacteria with native type V-K elements to ensure that these rare events are not missed. Alternatively, the BCQ pathway may be an evolutionary relic of a primitive selfish pathway before these transposons acquired CRISPR-Cas-based targeting modules.

From a technology perspective, type V-K CASTs are among the most compact type of CRISPR-associated transposases, in terms of coding size, and thus offer a major potential opportunity relative to type I CAST systems. However, two key properties that limit their use for genome engineering applications are low on-target specificity and the generation of cointegrate transposition products due to lack of TnsA^14, 15, 18, 45^. Recent efforts substantially decreased cointegrate formation by fusing an endonuclease to TnsB, and improved specificity using chimeric fusion proteins or supplementing additional components like pir^20^. Yet these strategies also compromise on-target integration efficiency and do not address the root cause of promiscuity. In this regard, our results provide a deeper molecular understanding of how type V-K CAST components undergo both RNA-guided and RNA-independent transposition. We identify TnsC filamentation on AT-rich DNA sequences as the primary driver of untargeted integration and show that, under limiting TnsC concentrations, RNA-guided transposition becomes the primary pathway of choice biochemically and in cells. This rescue in specificity was generalizable for all the guides we tested and importantly, resulted in equal or higher on-target integration efficiency when compared to the original ShCAST pHelper. Altogether, our combined use of biochemical, structural, and single-molecule experiments reveal the mechanistic intricacies associated with target site selection by type V-K CAST and offer new opportunities for targeted DNA integration applications.

## METHODS

### Plasmid construction

All constructs used in experiments with ShCAST were cloned using the pHelper (pUC19) and pDonor (pCDFDuet1) described previously^14^. These constructs were derived from a CAST system from *Scytonema hofmanni* UTEX B 2349^11^. Cloning was performed using a combination of Gibson assembly, inverse (around-the-horn) PCR, restriction digestion and ligation. All fragments used for cloning were amplified using Q5 DNA polymerase (NEB). TnsC and Cas12k were cloned into pCOLADuet1 vector backbone using Gibson assembly. The promoter strength was varied by cloning inducible promoters (lac, T7) or constitutive promoters (J23114, J23105, J23119) using inverse (around-the-horn) PCR. *S. hofmanni* S15 was cloned into expression strains and as fusions using a synthesized gene fragment (TWIST Biosciences). Different targets for pHelper or pHelper lacking TnsC was cloned in using around-the-horn PCR.

Constructs for protein expression having an N-terminal hexahistidine fused to a SUMO tag and a TEV cleavage site (His_6_-SUMO-TEV) were designed by restriction digestion of p1S vector (QB3 MacroLab) with Ssp1-HF(NEB) followed by Gibson assembly of the protein of interest. Expression was performed in *E. coli* BL21 (DE3) cells.

pTarget was designed by cloning a target from λ-DNA into pUC19 vector. Modulating regions in pTarget for introducing artificial untargeted sites was achieved using inverse (around-the-horn) PCR.

All primers and DNA oligos for the study were ordered from IDT. Cloning was performed in NEB Turbo *E. coli* and plasmid was extracted using Qiagen Miniprep columns and clones were confirmed by Sanger Sequencing (GENEWIZ). Whole plasmid sequence was confirmed by nanpore long read sequencing (Plasmidsaurus). *E. coli* was grown in LB agar plates or in liquid LB media. Constructs with pUC19 backbone was grown in 100 μg/mL carbenicillin, pCOLADuet1 and p1S in 50 μg/mL kanamycin and pCDFDuet1 in 100 μg/mL spectinomycin. All plasmid sequences and its description are available in **Supplementary Table 1**. Sequences of recombinantly expressed proteins used in this study is available in **Supplementary Table 2**.

### *E. coli* transposition assay

Transposition assay was performed in *E. coli* BL21(DE3) based on a method described previously ^14^, using a 2-plasmid system composed of pDonor, and pHelper. Chemically competent cells having pDonor were transformed with pHelper, plated, and grown at 37 °C for 16 h in the presence of appropriate antibiotics (100 μg/mL carbenicillin and 100 μg/mL spectinomycin). Cells were scrapped off and resuspended in LB. 4 x 10^9^ cells were re-plated, induced and grown in the presence 0.1 mM IPTG and antibiotics for 24 h at 37 °C. Cells were scrapped off and resuspended in LB. 2 x 10^9^ cells were used for genomic extraction using Wizard® Genomic DNA Purification

Kit (Promega). For transposition with an additional plasmid pCas12k, pTnsC or pTnsB, competent cells having pDonor, pCas12k/pTnsC/pTnsB were transformed with the pHelper and grown at the same conditions in the presence of appropriate antibiotics (50 μg/mL kanamycin, 100 μg/mL carbenicillin and 100 μg/mL spectinomycin). All *E. coli* transposition experiments were performed in n = 2 biological replicates.

### Recombinant expression of CAST proteins

All CAST proteins were cloned into a pET derivative vector with an N-terminal His_6_-SUMO-TEV fusion and were purified similar to previous protocols^11, 24^. All constructs were transformed in *E. coli* BL21(DE3) and grown until an OD600 of 0.5-0.6 in 2xYT media in the presence of kanamycin (50 μg/mL). Protein expression was induced in the presence of 0.5 M isopropyl ß-D-1-thiogalactopyranoside (IPTG) for 16 h at 16 °C.

In order to purify TnsB and TniQ, cells were harvested by centrifuging at 6000 g for 15 min at 4 °C and lysed by sonication in lysis buffer (20 mM Tris pH 7.5, 0.5 M NaCl, 5 mM Mg(OAc)_2_, 10 mM imidazole, 0.1% Triton X-100, 1 mM DTT and 5% (v/v) glycerol) supplemented with 1 mM phenylmethylsulfonyl fluoride (PMSF), 0.2 mg/mL lysozyme and 0.5x cOmplete protease inhibitor cocktail tablet (Roche). Soluble protein was isolated by centrifuging at 35,050g for 50 min and the supernatant was incubated with Ni-NTA agarose beads, pre-equilibrated with equilibration buffer (20 mM HEPES pH 7.5, 0.5 M NaCl, 0.5 mM PMSF, 1 mM DTT, 10 mM imidazole and 5% (v/v) glycerol) for 45 min at 4 °C. The Ni-NTA beads were further washed with 20 column volume (CV) of wash buffer (20 mM HEPES pH 7.5, 0.5 M NaCl, 0.5 mM PMSF, 1mM DTT, 30 mM imidazole and 5% glycerol) in a gravity column and protein was eluted in elution buffer (20 mM HEPES pH 7.5, 0.5 M NaCl, 0.5 mM PMSF, 1 mM DTT, 300 mM imidazole, and 5% glycerol). Next, His_6_-SUMO-tag was cleaved-off using 5% (w/v) TEV protease and was simultaneously dialyzed in Slide-a-Lyzer cassette (Thermo Fischer Scientific) against dialysis buffer (20 mM HEPES pH 7.5, 0.2 M NaCl, 0.5 mM PMSF, 1 mM DTT, and 1% (v/v) glycerol) for 12 h at 4 °C. Any precipitated protein was removed by centrifugation at 20,000 g for 20 min at 4 °C. Further, proteins were subjected to Heparin-affinity chromatography using a 0.2-1 M NaCl gradient spanning 10 CV in a buffer containing 20 mM HEPES pH 7.5, 0.2 M NaCl, 1 mM DTT and 1% glycerol. It is to be noted that TniQ does not bind to Heparin and elute in the flowthrough, whereas contaminant proteins were removed in this step. Further, both the proteins were purified by size-exclusion chromatography in SEC Buffer (20 mM HEPES pH 7.5, 0.2 M NaCl, 1mM DTT and 1% glycerol) using Superdex 200pg 16/600 column. TniQ and TnsB were concentrated to 127 and 100 µM and stored in a buffer containing 20 mM HEPES pH 7.5, 0.25 M NaCl, 5% glycerol, 1 mM DTT at -80 °C. We note no contamination of S15 in our TniQ potentially due the step involving Heparin (**Supplementary Table 2**)

For Cas12k, lysis, equilibration, wash, and elution buffer contained Tris pH 8.0 instead of HEPES pH 7.5. The dialysis buffer used contained 20 mM HEPES pH 7.5, 0.25 M NaCl, 0.5 mM PMSF, 1 mM DTT, and 1% (v/v) glycerol. The SEC buffer contained 20 mM HEPES pH 7.5, 0.25 M NaCl, 1 mM DTT, and 1% (v/v) glycerol. All the rest of the protocol and buffers used were the same as in case of TnsB and TniQ. Cas12k was concentrated to 109 μM and stored at the same conditions as TnsB and TniQ.

For TnsC, lysis, equilibration, wash, and elution buffer contained 0.5 M NaCl to increase protein solubility. TEV cleavage and dialysis was performed in 20 mM HEPES pH 7.5, 1 M NaCl, 0.5 mM PMSF, 1 mM DTT, and 1% (v/v) glycerol after which the protein was subject to a second round of Ni-NTA to separate His_6_-SUMO tag from the cleaved TnsC. The column was washed with 20 mM HEPES pH 7.5, 500 mM NaCl, 30 mM imidazole, 1 mM DTT, 0.5 mM PMSF, 5% (v/v) glycerol, 1 mM DTT, 0.5 mM PMSF and the flowthrough was concentrated and further purified by size exclusion chromatography in 20 mM HEPES pH 7.5, 1 M NaCl, 5% glycerol, 1 mM DTT using Superdex 200pg 16/600 column. TnsC was concentrated to 142 µM and stored in a buffer containing 20 mM HEPES pH 7.5, 1 M NaCl, 10% (v/v) glycerol, 1 mM DTT.

For S15, lysis, equilibration, wash, and elution buffer contained Tris pH 8.0 instead of HEPES pH 7.5. The dialysis buffer contained 20 mM HEPES pH 7.5, 0.15 M NaCl, 0.5 mM PMSF, 1 mM DTT, and 1% (v/v) glycerol. S15 was eluted from Heparin using a gradient of 0.15 to 1 M NaCl in 20 mM HEPES pH 7.5, 1mM DTT and 15% (v/v) glycerol. SEC contained 20 mM HEPES pH 7.5, 0.25 M NaCl, 1mM DTT, and 1% (v/v) glycerol. All the rest of the protocols and buffers used were the same as in case of TnsB and TniQ. S15 was concentrated to 168 µM and stored at the same conditions as TnsB and TniQ.

### Untargeted transpososome sample preparation

The untargeted transpososome was reconstituted as previously described for the Cas12k-bound transpososome^26^, with some modifications noted below. Two separate DNA substrates were first prepared by annealing three synthetic oligonucleotides (IDT) respectively: LE1, LE2, and LE3 for target-LE DNA and RE1, RE2, and RE3 for target-RE DNA. The oligonucleotides were mixed in a 1:1:1 molar ratio and supplemented with 10X concentrated annealing buffer for the following final composition: 10 mM Tris pH 7.5, 50 mM NaCl, and 1 mM EDTA. These mixtures were then heated up to 95 °C for 5 minutes and slowly cooled to room temperature at the rate of 1 °C per minute using a thermal cycler (BioRad).

TnsB, TnsC, and TniQ proteins were purified identically as previously described^11, 26^. All the protein stocks were first buffer-exchanged into the dilution buffer (25 mM HEPES pH 7.5, 200 mM NaCl, 1 mM DTT, and 2% glycerol) using 0.5 mL centrifugal filters (Millipore). For the target pot, TnsC, TniQ, and the annealed target-LE DNA were first mixed and supplemented with ATP and MgCl_2_, resulting in the final composition of the following: 45 µM TnsC, 30 µM TniQ, 3 µM target-LE DNA, 3 mM ATP, and 10 mM MgCl_2_ in the dilution buffer. For the donor pot, TnsB and target-RE DNA were combined to make the following composition: 36 µM TnsB, 6 µM target-RE DNA, and 10 mM MgCl_2_ in dilution buffer. Target and donor pots were incubated independently for 10 min at room temperature before being combined in a 2:1 volume ratio to reconstitute the transpososome. The composition of the final reaction condition was the following: 30 µM TnsC, 20 µM TniQ, 12 µM TnsB, 2 µM target-LE DNA, 2 µM target-RE DNA, 2 mM ATP, and 10 mM MgCl_2_. The final reaction mixture was then incubated for 30 min at 37 °C.

### Cryo-EM sample preparation and imaging

Homemade graphene-oxide (GO) coated grids were made as previously described^35, 46^ or cryo-EM sample preparation. The untargeted transpososome sample was prepared by diluting the final reconstitution mixture three-fold using the dilution buffer (described above). Four microliters of the diluted sample were loaded on the carbon side of the GO-coated grid, which was mounted on the Mark IV Vitrobot (ThermoFisher) set to 4 °C and 100% humidity. The grid was then incubated for 20 seconds in the Vitrobot chamber to let the proteins adhere to the GO surface of the grid. Then the grid was blotted for 7 seconds with a blot force of 5 before being plunged into a slurry of liquid ethane cooled with liquid nitrogen.

The vitrified samples were imaged using 200 kV Talos Arctica (ThermoFisher) equipped with K3 direct electron detector (Gatan) and BioQuantum energy filter (Gatan). The electron beam was carefully aligned following the established protocol^47^ or parallel illumination and coma-free alignment. 3,081 micrographs were recorded using SerialEM^48^ at 63,000X magnification, corresponding to 1.33Å per pixel scale. A three-by-three image shift was used to accelerate the image collection with a nominal defocus from -1 µm to -2.5 µm. The total electron dose was 50 electrons per 1Å^2^ during 3.2 seconds of recording, fractionated into 50 frames.

### Cryo-EM image analysis and visualization

Warp^49^ as used for beam-induced motion correction, CTF estimation, and initial particle picking of the collected movies. The preprocessed micrographs were filtered based on the CTF-estimated resolution, resulting in 2,790 micrographs. These micrographs and corresponding particle stacks were imported to cryoSPARC^50^ or downstream image analysis. 2D classification of the particle stack resulted in a subset of particles with a cryo-EM density of both TnsC oligomer and TnsB strand-transfer complex (STC). This set of particles was used to train the Topaz^51^ neural network, which extracted 993,190 particles. 2D classification of this initial particle stack resulted in 577,628 particles with a cryo-EM density of TnsB STC and TnsC. These particles were then subjected to heterogeneous refinement in cryoSPARC using the three initial references of the following: (1) TnsB STC only, (2) two turns of TnsC, and (3) TnsB STC bound to two turns of TnsC, which was generated using ab initio reconstruction. 237,487 particles classified into the third class were then subjected to focused refinement on the TnsC region using a mask covering the TnsC. The resulting particles were then exported to RELION^52^ or 3D classification focusing on TniQ and adjacent two TnsC subunits. Since the molecular mass of this region was relatively small (∼80 kDa), 3D classification was done without image re-alignment. One class (42,210 particles), corresponding to strong density of TniQ, was selected and re-imported into cryoSPARC for another round of 3D classification, focusing on the single subunit of TniQ. This final classification resulted in a final stack of 13,392 particles. This final particle stack was subjected to homogeneous refinement to generate a consensus map. Focused refinements were done through two separate non-uniform refinement jobs focusing on the TniQ-TnsC region or the TnsB STC region of the reconstruction. Local resolution and directional resolution were estimated using blocres^53^ and 3DFSC^54^, respectively. For **Fig. 1f** local-resolution filtered maps from each focused refinement were aligned onto the consensus map and then combined using UCSF Chimera^55^ command ‘vop maximum’. Atomic model was generated through rigid-body docking of the chains from the RNA-guided transpososome structure (PDB: 8EA3)^23^. Figures describing the cryo-EM reconstruction or atomic model were generated using UCSF ChimeraX^55^.

### *E. coli* growth time course

*E. coli* Bl21(DE3) cells were transformed with a plasmid containing combinations of TnsC and TnsB on an inducible promoter (T7) or an empty plasmid and grown for 16 h in the presence of appropriate antibiotics (50 μg/mL kanamycin and 100 μg/mL carbenicillin). Single colonies were picked for each biological replicate and inoculated to a primary culture and grown for 16 h with antibiotics. 1:200 dilution of primary culture was added to sterile 96-well plate black/clear bottom plates (Thermo Scientific) in a final volume of 200 µL with 0.1 mM IPTG and antibiotics. The culture was grown at 37 °C with continuous shaking and OD600 was measured for 16 h in Synergy Neo2 plate reader (BioTek). All readings were blank corrected. The error bar represents the standard deviation measured across a minimum of two biological replicates.

### Fluorescence polarization

Fluorescence polarization was performed as reported earlier^33^. A 55 bp dsDNA labeled with a 5’ Fluorescein (6-FAM) tag on one of its strands was used for all fluorescence polarization experiments (**Supplementary Table 3**). Recombinantly expressed TnsC (0-15 μM) was titrated with the FAM-labeled dsDNA in 1X binding buffer (20 mM HEPES pH 7.5, 2 mM MgCl_2_, 200 mM NaCl, 10 µM ZnCl_2_, 1 mM DTT) in the presence of 2 mM ATP or its analogs. The samples were incubated at 37 °C for 30 min in a 384-well plate. The fluorescence gain was adjusted to a sample lacking TnsC while keeping a requested polarization value of 60 mP. The error bar represents the standard deviation measured across two technical replicates. For low salt measurements, the concentration of NaCl was maintained at 50 mM.

For fluorescence polarization measurements for TniQ alone or its interaction with DNA bound to TnsC, TniQ (0-1 µM) was titrated and incubated for 30 min at 37 °C with fluorescently labeled DNA alone or DNA preincubated with 300 mM TnsC (30 min at 37 °C). Experiments were conducted in a 1X binding buffer containing 50 mM NaCl in a 25 µL final volume.

### *In vitro* ATP hydrolysis assay

Relative ATP hydrolysis was measured using a Malachite Green Phosphate Assay Kit (Sigma-Aldrich) according to the manufacturer’s protocol as previously described^33^. TnsC (10 µM) alone or TnsC (10 µM) together with TnsB (20 µM) was incubated with 10 µM dsDNA (**Supplementary Table 3**) in the presence of 1X ATPase reaction buffer (20 mM HEPES pH 7.5, 2 mM MgCl_2_, 180 mM NaCl, 10 μM ZnCl_2_, 1 mM DTT) supplemented with 1 mM ATP in a final 5 µL volume for 120 min at 37 °C. The ATP hydrolysis reaction was diluted with 40 µL water and incubated with 10 µL of working reagent for 5 min. OD at 620 nm was measured in a transparent 384-well plate in plate reader. All samples were blank corrected. The ATPase activity measured was normalized to the sample containing TnsC, TnsB and dsDNA which liberated 174 µM of phosphate. Error bar represents the s.d. measured across two technical replicates (n = 2).

### Biochemical transposition assay

Guide RNA was transcribed *in vitro* using MEGAclear Transcription Clean-Up Kit (Invitrogen) using a linear dsDNA as a template (**Supplementary Table 3**). Prior to setting the transposition reaction, the ribonucleoprotein (RNP) complex composed of Cas12k (0.75 μM) and guide RNA (sgRNA-6, 9 μM) was pre-incubation at R.T. for 5 min in a total volume of 1.4 µL. *In vitro* integration was performed in the presence of RNP (50 nM), TniQ (100 nM), TnsC (100 nM), TnsB (1 μM), pTarget (0.6 nM) and pDonor (1.75 nM) supplemented with ATP (2 mM), BSA (50 μg/mL), SUPERase•In Rnase Inhibitor (Invitrogen) (0.25 U/μL) in a 1X IVI Buffer (25 mM HEPES pH 7.5, 5 mM Tris pH 8.0, 0.05 mM EDTA, 20 mM MgCl_2_, 30 mM NaCl, 20 mM KCl, 1 mM DTT, 5% (v/v) glycerol). The reaction having a final volume 25 µL was incubated at 37 °C for 2 h, quenched by heat denaturing at 95 °C for 3 min and flash frozen until library preparation for NGS. Transposition was performed in the presence of Mg^2+^ ion concentration of 2 mM throughout the reaction, in contrast to the conditions mentioned earlier^11, 56^. We note that pre-incubation with low Mg^2+^ at 30 °C is not necessary for the biochemical reconstitution of CAST transposition. For titration involving individual proteins, the concentration was varied as indicated (**Fig. 2**, **Supplementary Fig. 4**), keeping all other components constant. Indicated concentrations of S15 was supplemented for reactions involving the same. The product of *in vitro* transposition are Shapiro intermediates as opposed to resolved co-integrates in cellular experiments. The transposon-target DNA junction corresponding to the Shapiro intermediate was amplified by PCR and visualized on 1.5% agarose. qPCR, or NGS library preparation was also performed on the Shapiro intermediate. Due to linear amplification in the first cycle of PCR for the Shapiro intermediate compared to the reference used in qPCR, the efficiencies measured are relative for biochemical experiments. Without S15, we observed a relative on-target efficiency of ∼10% detected by qPCR whereas supplementing S15 resulted in 33% efficiency.

### qPCR analysis for *in vitro* and genomic transpositions

qPCR for detecting on-target integration in the genome was performed with two sets of primers. A combination of – 1) genome-specific (targets for sgRNA 1-5) and left transposon-specific primers probing t-LR and 2) genome-specific primers for detecting an *E. coli* reference gene *rssA*. Primer pairs were designed to amplify products between 100 and 250 bp and showed amplification efficiency between 100 and 110 %. Probes for Taqman qPCR were designed with a 5’ FAM-label ZEN/3’ IBFQ (IDT) DNA as a universal probe which annealed to the transposon left-end and can be displaced only upon amplification. The control probe was designed to bind to *E. coli rssA* gene and contained a 5’ SUN-label ZEN/3’ IBFQ (IDT) DNA. qPCR for genomic samples was performed using 5 ng of purified genomic DNA in the presence of 1 µL each of 18 µM forward primer and reverse primer pairs (**Supplementary Table 3**), 5 µL of TaqMan Fast Advanced Master Mix (Thermo Fischer Scientific), 0.5 µL of each 5 µM probe (**Supplementary Table 3**) and 1 µL of water in a final volume of 10 µL. On-target integration efficiency is measured as 100 X (2^ΔCq), where ΔCq is the Cq (control rssA) – Cq (integration).

qPCR for detecting the Shapiro intermediate resulting from on-target integration was performed with two sets of primers. A combination of – 1) pTarget and left transposon-specific primers probing t-LR and 2) primers for detecting pTarget. qPCR measurements for *in vitro* integration were performed with 2 µL of a 20 µL reaction, 1 µL each of 10 <u>µM</u> forward and reverse primer (**Supplementary Table 3**), 5 µL of SsoAdvanced Universal SYBR Green 2X Supermix (BioRad), and 2 µL of water in a final volume of 10µL. The first cycle of qPCR with primers detecting on-target integration only results in linear amplification at the Shapiro intermediate in comparison to the control primers detecting pTarget and therefore we note that the efficiency measurements are only relative. On-target integration efficiency is measured as 100 X (2^ΔCq), where ΔCq is the Cq (pTarget) – Cq (integration).

All qPCR reactions were performed on 384-well clear/white plates (BioRad) on a CFX384 Real-Time PCR Detection System (BioRad) using the following thermal cycling conditions: DNA denaturing (DNA denaturation 98 °C for 2.5 min), 40 cycles of amplification (98 °C for 10 s, 62 °C for 20 s), and terminal melt-curve analysis was performed (65–95 °C in 0.5 °C per 5 s increments). All qPCR primers used in this study are mentioned in **Supplementary Table 3**.

### TagTn Seq library preparation and sequencing

In order to get usable sequencing coverage across genomic and plasmid targets, samples from biochemical and *E. coli* transposition assays were treated with AvrII (NEB) to reduce the proportion of contaminating pDonor. A single AvrII cut site was located 21-nt from the left transposon-end in pDonor whereas pTarget had no sites. *E. coli* BL21-DE3 genome contains only 16 AvrII cut sites. For *in vitro* integration, 1.5X µL of Mag-Bind TotalPure NGS magnetic beads (Omega) were added to each sample and the DNA was purified using the manufacturer’s protocol. The sample was eluted in 10 µL volume and digested with AvrII (NEB) (5 U) in rCutSmart Buffer (NEB) for 1 h at 37 °C. 1.5X magnetic beads were added to each sample and DNA was purified and eluted in a 10 µL volume. For genomic samples, 1 µg of isolated genomic DNA (having contaminating pDonor) was digested with AvrII (5 U) in a 50 µL volume for 2 h at 37 °C. Total genomic DNA was purified with magnetic beads, eluted in 10 µL volume and quantified using Qubit dsDNA High Sensitivity Kit (Invitrogen) and was used for tagmentation using Nextera XT DNA Library Preparation Kit (Illumina).

4 ng of DNA from either biochemical or *E. coli* transposition assays were mixed with 5 µL of tagmentation DNA buffer (Illumina) and 1 µL of amplicon tagmentation mix (Illumina) in a final volume of 10 µL and incubated at 55 °C for 7 min. The tagmentation reaction was quenched and DNA was amplified by PCR-1 step, using CAST left transposon-end specific primer, Y3 mix (0.42 µM), having universal TrueSeq adaptor (i5) overhang, Nextera adaptor (i7) specific primer (0.42 µM) (**Supplementary Table 3**) and 30 µL KAPA HiFi HotStart ReadyMix in a 60 µL final volume. Y3 mix is a combination of three primers with variable length between CAST transposon-end specific region and universal TrueSeq adaptor to introduce diversity during Illumina flow cell clustering (**Supplementary Table 3**). After 20 cycles of amplification at an annealing temperature of 57 °C, the amplified DNA was purified using 1.5X of magnetic beads and eluted in 10 µL volume. This DNA was next subjected to PCR-2 using TrueSeq (i5) specific primers (0.42 µM) having indexed universal p5 overhang, Nextera adaptor (i7) specific primer (0.42 µM) having indexed universal p7 overhang (**Supplementary Table 3**), and 30 µL KAPA HiFi HotStart ReadyMix in a 60 µL final volume. After 13 cycles of amplification at an annealing temperature of 54 °C, the barcoded amplicon was purified by magnetic beads and eluted in 10 µL volume. The DNA was resolved by 1.5 % agarose followed by excision and isolation within the size range of 300-600 bp using Gel Extraction Kit (Qiagen). NGS libraries were pooled and quantified by qPCR using NEBNext Library Quant Kit (NEB). Sequencing was performed with NextSeq high-output kit with 75/150-cycle single-end reads and automated adaptor trimming and demultiplexing (Illumina) was performed.

### Analysis of NGS data

Analysis of TagTn Seq data was performed using a custom Python pipeline as described^14^. Demultiplexed raw reads having half of the bases with a Phred quality score of less than 20, which corresponds to greater than 1% base miscalling were removed from the analysis. Reads having the last 23 bases of the transposon-end sequence (5’ GACAGATAATTTGTCACTGTACA 3’) and a 22-bp adjacent flanking genomic sequences were extracted and noted as the total-transposon-end containing reads. This 22-bp adjacent region represents the fingerprint region used to identify transposon insertions and was used to align to the reference genome or plasmid using Bowtie2^57^. The reference genome for *E. coli* BL21(DE3) was based on published data from National Center for Biotechnology Information (NCBI) genomes. Only reads which mapped perfectly and only once to the genome were chosen. Reads which did not get mapped to the genome were checked for sequences corresponding to the pDonor and noted as pDonor contamination. Alignments from Bowtie2 were used for generating genome/plasmid-wide coordinates for integration. If the read was mapped to the same strand as the input fasta file, then it was noted as on ‘fwd’ strand, and if mapped to the complementary strand as the input fasta file, then it is noted as on ‘rev’ strand. The read ‘position’ was indexed at the fifth position of target site duplication (TSD) for each event, with respect to the ‘fwd strand’. The orientation of the integration relative to the fasta file was concluded based on whether the library was sequenced from the right end or the left end of the transposon for TagTn sequencing. The orientation of the transposon insertion with respect to the protospacer at the on-target window (100 bp from the end of protospacer), was noted as target-left-right (t-LR) or target-right-left (tRL).

Untargeted reads and on-target reads were assigned using a custom Python script. For biochemical integration, events that mapped at a hundred base window after the end of the protospacer on pTarget were presumed as on-target events, whereas any other integration events on pDonor and pTarget were totaled and noted as untargeted. Reads due to contaminating pDonor and a potential PCR recombination artifact (at position 1198 and 2328 nt) were masked for all samples. For *E. coli* integration, reads that mapped at a hundred base window after the end of the protospacer were noted as on-targets, and reads elsewhere in the genome as untargeted. Raw reads for integration at on-target and untargeted sites for each sample was normalized with the total transposon-end containing reads (including pDonor contamination) detected for that sample and further scaled with respect to the sample having the highest transposon-end containing reads (in the experimental set). It is to be noted that we do not correct for amplification biases during NGS sequencing and for this reason, only unique insertion sites were used for reporting sequence preferences in transposition. For visualizing normalized plots comparing integration across the genome, raw reads for each coordinate for a sample were normalized and scaled in the same way as above, converted to bigwig files and visualized in IGV.

For the plot comparing untargeted integration sites across pTarget, reads at the on-target window were excluded. A continuous single-base position for the pTarget and the reads detected at that position was generated using a custom Python script. The read at each of these bases were compared between samples. Pearson correlation coefficients (r) between data sets were calculated in Prism 9.

### Analysis of genetic neighborhood for integration events detected across the *E. coli* genome

A custom Python script was used to analyze whether the integration events detected elsewhere in the genome are RNA-dependent off-targets. The position of insertion determined by NGS analysis was used for the genetic neighborhood analysis. Events at the on-target window were removed from the analysis. For every integration site detected, the mostly likely RNA-dependent off-target was determined as follows. Off-target was assumed to be in the target-left-right orientation similar to an on-target event. A 7 bp window (3 bp upstream and downstream) of sequence, 63 bp upstream from the integration site in the LR orientation was checked for the enrichment of a PAM sequence (GTN). Similarly, n = 10,000 random regions in the *E. coli* genome were chosen, and the corresponding window upstream was checked to get the proportion of random sequences having a potential PAM.

For integration events, PAM-containing sequences were further checked for the presence of matches in the protospacer for sgRNA 1-5 respectively (23 bp downstream to the potential PAM detected). The maximum matching sequence detected was selected as the potential number of spacer matches representing that insertion event. Similarly, the random region of *E. coli* genome sampled previously with a PAM-containing sequence was checked for matches in protospacer for sgRNA 1-5 respectively. In both cases, when more than one potential protospacer had a valid PAM, the maximum number of matches to the spacer was determined as the potential number of spacer matches for that location.

### AT-enrichment analysis

AT-enrichment analysis was performed with a custom Python script. The position of insertion determined by NGS analysis was used for analyzing the AT content. Events at the on-target location were removed from the analysis if a spacer was provided. For every unique insertion site on pTarget, λ-DNA or the *E. coli* genome, a window of sequence either upstream or downstream was chosen (50 bp for pTarget or λ-DNA and 100 bp window for the *E. coli* genome). The AT content at both these regions (upstream/downstream) was calculated and the highest of the two was selected as the representative AT content of that unique insertion site. Unique events were binned to across their AT content to create a distribution of events detected at each AT bin. As a control to calculate the AT content of pTarget, λ-DNA or the *E. coli* genome, the regions on the DNA were randomly sampled (n = 50,000) and similarly, the highest AT content of the adjacent bin to every unique sampled location was chosen as representative of that sampling event. The randomly sampled events were also binned across their AT content to create a distribution of random samplings events (counts) detected at each AT bin. A cumulative frequency distribution plot was used to compare the difference in AT content for unique integration events and randomly sampled regions on the pTarget, λ-DNA or the *E. coli* genome. The AT content of unique integration events and random sampling events were used to perform Mann-Whitney U test used to test the significance of the distributions. pDonor was not used to for AT-enrichment analysis due to additional factors such as target immunity^58^.

To generate plots to visualize the AT content across the pTarget and integration, the pTarget was divided into 59 bins corresponding to 46 bp and the AT percentage was calculated using a custom Python script. The reads detected on pTarget was plotted as an overlay with AT content at that bin. For λ-DNA, the genome was into 45 bins of 1078 bp window and AT percentages were calculated. The integration events detected in λ-DNA were plotted as an overlay with the AT content. Correlation between reads detected for integration and fluorescence intensity of mNG-TnsC binding across λ-DNA (from DNA curtains) were calculated by similarly binning the reads and fluorescence intensity detected across λ-DNA into 45 bins of 1078 bp window and Spearman correlation coefficient (r) was calculated using Prism 9.

### Essential gene analysis

Essential gene analysis (**Supplementary Fig. 5l**) was performed with a custom Python script. The position of insertion determined by NGS analysis was used to determine if events landed on an essential gene or not. Events at the on-target location and at the T7RNAP locus were removed from the analysis. The remaining reads for these events were annotated with their CDS features in accordance with the NCBI-published genome. Essential genes were noted based on previous reports on *E. coli* K-12 and reads falling to these regions were classified as essential gene insertions^59^. Reads landing outside these regions were noted as non-essential insertions. The percentage of the *E. coli* genome which was essential was calculated by summing the length of all essential genes and dividing by the genome length.

### Sequence logo for untargeted events

To create sequence logos for untargeted events, we took every unique site of integration in the *E. coli* genome, and a window upstream and downstream to the position of integration on both the ‘fwd’ and ‘rev’ strand. Insertions in the on-target window were omitted for samples having a spacer. Further, TSD correction was accounted on the strands, and insertions were oriented in the left-to-right orientation based on the strand of the input fasta file and the strand to which each read was mapped. The extracted sequence for both the strands was outputted and WebLogo v2.8.2 was used for plotting the sequence logo showing per residue conservation.

For ShCAST, a 75-base window on either side of integration was chosen for building the sequence logo. The sequence logo was indexed with the third position of TSD set as position ‘0’ and 70 bases on either side was plotted. WebLogo^60^ in **Fig. 3h** was plotted for insertions on both the strands observed for two biological replicates of a sample having a pHelper lacking Cas12k and guide RNA. Sequence features for samples with Cas12k and guide RNA (sgRNA-1) was performed using previously reported data for ShCAST^14^ generated by random fragmentation and insertion site amplified from the transposon right-end (**Supplementary Fig. 6b**). The plots revealed consistent sequence featured compared to samples subjected to TagTn Seq and sequencing from the transposon left end. No conservation of sequences with partial guide complementarity was noted upstream, consistent with the analysis (**Supplementary Fig. 1c-e**) that the majority of events in type V-K CASTs are RNA-independent.

For ShoCAST, previously reported data^14^ with a pHelper having ShoCas12k and guide RNA (sgRNA-7) was used and analyzed to inform the position and read for each integration event. WebLogo^60^ in **Supplementary Fig. 6c** was plotted for unique insertions. A 75-base window was chosen for building the sequence logo. The sequence logo was indexed with the third position of TSD set as position ‘0’ and 70 bases on either side were plotted.

### DNA curtains

Single-molecule double-tethered dsDNA curtain experiments were carried out as previously described^36^. Custom flow cells were assembled using quartz slides, on which chromium patterns were deposited through nanofabrication^61^. The dsDNA substrate was prepared by annealing and ligating two custom oligonucleotide handles (IDT) at the COS sites of bacteriophage λ-DNA (NEB), such that one end of the DNA contained a biotin modification while the other a digoxigenin (dig). Prior to assembly of DNA curtains, flow cells were first passivated with a lipid bilayer. The biotin ends of the DNA were tethered through a biotin-streptavidin linkage to biotin-modified lipids within the bilayer. These DNA molecules were then aligned as the chromium barriers and flow-stretched so that the dig-tagged ends were anchored to chromium pedestals coated with anti-dig antibody (Roche). These DNA molecules were then flow-stretched so that the dig-ends were anchored at chromium pedestals coated with anti-dig antibody (Roche). A pre-defined barrier-pedestal distance of 12 µm and orthogonal attachment chemistry yielded uniform and unambiguous orientation of the double-tethered dsDNA curtain. Fluorescent mNeonGreen-TnsC (mNG-TnsC) was injected into the flow cell, from a 50 µL sample loop, using a syringe pump. Sample illumination was achieved using a continuous-wave 488 nm laser (Coherent Sapphire, 200 mW), shuttered externally (Uniblitz LS6). The emission signal was visualized through a custom-built prism-type total internal reflection fluorescence (TIRF) microscopy system, based on a Nikon Eclipse TE2000-U, equipped with a 60X water immersion objective and an EMCCD camera (Andor iXon X3). All single-molecule experiments were performed in reaction buffer (20 mM HEPES pH 7.5, 2 mM MgCl_2_, 200 mM NaCl, 10 µM ZnCl_2_, 1 mM DTT, 0.2 mg/mL BSA, and 2 mM ATP), at room temperature (25 °C). In equilibrium binding experiments, flow was stopped after mNG-TnsC (100 nM) had reached the flow cell and data were collected at the rate of 1 frame per 30 seconds for 30 min. mNG-TnsC was injected at a concentration of 500 nM for the experiment shown in **Supplementary Fig. 5c** and **Supplementary Movie 2**.

### Data analysis for DNA curtains

Files saved in ND2 format were first converted to TIFF stacks in FIJI^62^, where all subsequent image processing steps took place. For binding and disassembly experiments, a kymograph was first generated for each DNA molecule by making a single-pixel wide slice, drawn along the length of the DNA, through the time series. Fluorescence intensity over time was then extracted by plotting the profile of the kymograph of interest in FIJI, which produced intensity values, each averaged over the entire one-pixel-wide DNA, over time. Intensity time series from >50 individual filaments were combined, normalized, and plotted as mean ± s.d. over time (**Fig. 3c and 3d**). Apparent rates (*k*_obs_; **Fig. 3d**) were obtained by nonlinear regression to a one-phase association model in Prism 9. For intensity versus binding position (**Fig. 3c**), the last 10 frames (5 min) of a 30-minute binding experiment were averaged to remove intensity fluctuations in any given frame due to random DNA fluctuations within the evanescent field. For each DNA of interest, a signal intensity profile was plotted over a 33-pixel long region, centered over the 45-pixel total length of the DNA (12 µm), to reduce signal contamination and bleed-over from nonspecific binding of mNG-TnsC to chromium features. Intensity profiles from > 60 individual DNA molecules were combined and plotted as mean ± standard deviation. The A/T percentage of the λ-sequence was obtained by first segregating the entire sequence (48,502 bp) into 45 bins of 1078 bp width, then calculating the A/T percentage for the sequence within each bin. The number of bins was chosen to reflect the fact that λ-DNA double tethered at 12 µm end-to-end distance spanned 45 pixels in the camera of this imaging platform. Thus, fluorescence intensity measured at any pixel along the entire filament should represent the amount of mNG-TnsC bound to the DNA within the window of 1078 bp.

### Western blot

An N-terminal Flag epitope tag was cloned into TnsC expressed with a T7 or lac promoter. Cells were grown at the same conditions as the *E. coli* transposition assay in the presence of 0.1 mM IPTG. 2 X 10^7^ were incubated with 5 µL of 6X SDS loading dye and boiled for 10 min at 95 °C. The samples were resolved on Mini-PROTEAN TGX Stain-Free Precast Gels and imaged to confirm equal loading. The gel was transferred using iBlot 2 Gel Transfer Device using the manufacturer’s protocol. The transferred blots were washed twice with wash buffer (1% PBS, 0.1% Tween) for 5 min each and then washed with blocking buffer (1% PBS, 0.1% Tween, 5% BSA) for another 5 min. The blot was then incubated for 30 min in 70 mL blocking buffer at RT with gentle shaking. After 30 min, the blot was again washed with block buffer and incubated for 1h with either Monoclonal ANTI-FLAG M2 (Sigma) (1:7500 dilution) in 15 mL volume or GAPDH Loading Control Monoclonal Antibody (Thermo Scientific) (1:2500 dilution) in 5 mL volume in blocking buffer with gentle shaking. The same wash protocol was repeated and incubated with a secondary HRP conjugated antibody, Goat Anti-Mouse IgG1 HRP (Abcam) (1:15000 dilution) in 15 mL volume and incubated for 2 h. The blot was washed twice in the wash buffer for 5 min each and imaged using SuperSignal West Dura Extended Duration Substrate (Thermo Scientific) after incubating for 2 min.

### Statistics and reproducibility

All *E. coli* transposition assays and NGS were performed in two independent biological replicates. qPCR for the same was performed in two independent replicates. *E.* coli growth curves were measure in minimum of two independent replicates. Titration for *in vitro* transposition were performed once for each concentration. Single molecule experiments were done in replicates (data not shown). Fluorescence polarization and ATP hydrolysis assays were performed in two technical replicates.

### Data availability

Next-generation sequencing data will be made available in the National Center for Biotechnology Information (NCBI) Sequence Read Archive. Datasets generated and analyzed in the current study are available from the corresponding author upon reasonable request.

### Code availability

Custom python scripts used for computational analysis of next generation sequencing data are available at GitHub (https://github.com/sternberglab/George_et_al_2023).

## SUPPLEMENTARY INFORMATION

A list of all plasmids, purified proteins, primers, and guide RNAs used in this study is available in Supplementary Tables 1-4.

## ACKNOWLEDGEMENTS

We thank N. Jaber and S.R. Pesari for laboratory support, P.A. Sims for helpful discussions on deep sequencing, M. Jovanovic for help with mass spectrometry, R.T. King for help with Taqman qPCR, D.R. Gelsinger for help with TagTn Seq, G.D. Lampe, F.T. Hoffmann, and S. Tang for critical feedback on the manuscript, L.F. Landweber for qPCR instrument access, the JP Sulzberger Columbia Genome Center for NGS support and Cornell Center for Materials Research facility, K. Spoth, M. Silvestry-Ramos, for maintenance of electron microscopes used for this research (NSF MRSEC program, DMR-1719875). J.T.G. is supported by HFSP postdoctoral fellowship (LT001117/2021-C) from the International Human Frontier Science Program Organization. E.C.G. is supported by NIH grant R35GM118026. E.H.K. is supported by NIH grant R01GM144566 and a Pew Biomedical Scholarship. S.H.S. is supported by NIH grants DP2HG011650, R21AI68976, and R01EB031935, a Pew Biomedical Scholarship, a Sloan Research Fellowship, an Irma T. Hirschl Career Scientist Award, and a generous startup package from the Columbia University Irving Medical Center Dean’s Office and the Vagelos Precision Medicine Fund.

## AUTHOR CONTRIBUTIONS

J.T.G. and S.H.S. conceived of and designed the project. J.T.G. performed most experiments in the study. C.A. assisted in the analyses of high-throughput sequencing data and contributed computational support. J.P. and E.H.K. performed cryoEM experiments and data analysis. M.K. and E.C.G. assisted with single-molecule biophysics experiments. T.W. contributed bioinformatics and structural analyses. Y.L.P. assisted with protein biochemistry. J.T.G. and S.H.S. discussed the data and wrote the manuscript, with input from all authors.

## SUPPLEMENTARY FIGURES

**Supplementary Fig. 1.**
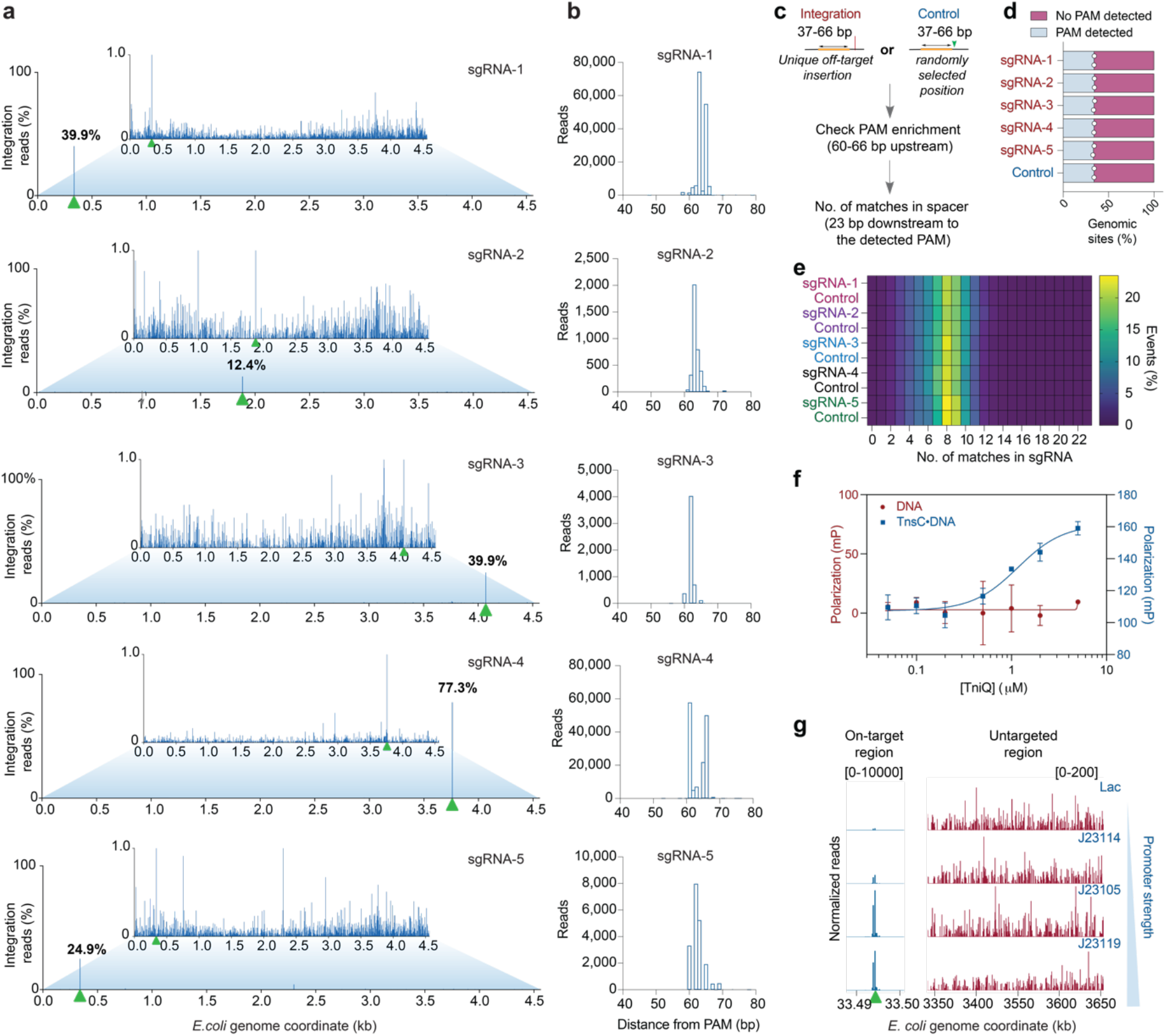
Type V-K CASTs maintain a distinct, RNA-independent pathway. **a**, Genome-wide view of *E. coli* genome-mapping reads for the original/WT ShCAST system encoding the indicated sgRNAs; the zoomed-in views visualize reads comprising 1% or less of genome-mapping reads. Target sites are marked with a green triangle. **b**, Integration site distributions determined from the NGS data, plotted as the distance from the PAM sequence to the first transposon bp. **c,** Schematic depicting the neighborhood analysis used to explore PAM (5’- GTN-3’) and sgRNA complementarity enrichment flanking off-target ShCAST integration events. Insertions were presumed to occur in the (target-left-right) T-LR^14^, and a 60-66 bp window upstream of the insertion site was analyzed. For sites with a cognate PAM, the adjacent 23 bp were analyzed for complementarity to the sgRNA spacer sequence. Controls analyses used random samplings of the *E. coli* genome. **d,** Off-target ShCAST integration events from five distinct sgRNAs are not enriched in cognate PAMs, relative to a randomly sampled control dataset (**Methods**). Percentage of integration events (for n = 2 biological replicates) detected with or without a PAM when compared to random regions of the *E. coli* genome (Methods). **e,** Off-target ShCAST integration events from five distinct sgRNAs are not enriched in the number of complementary matches to the sgRNA, relative to a randomly sampled control datasets (**Methods**). **f**, TniQ selectively binds TnsC-DNA filaments, but not naked DNA, as observed using fluorescence polarization experiments with a 55-bp FAM-labeled DNA substrate. Data shown represent mean ± s.d for n = 2 technical replicates. **g,** Normalized integration reads detected at the on-target site (green arrow, left) and a representative untargeted site (right), with varying Cas12k promoter strength. Note the differing y-axis ranges.

**Supplementary Fig. 2.**
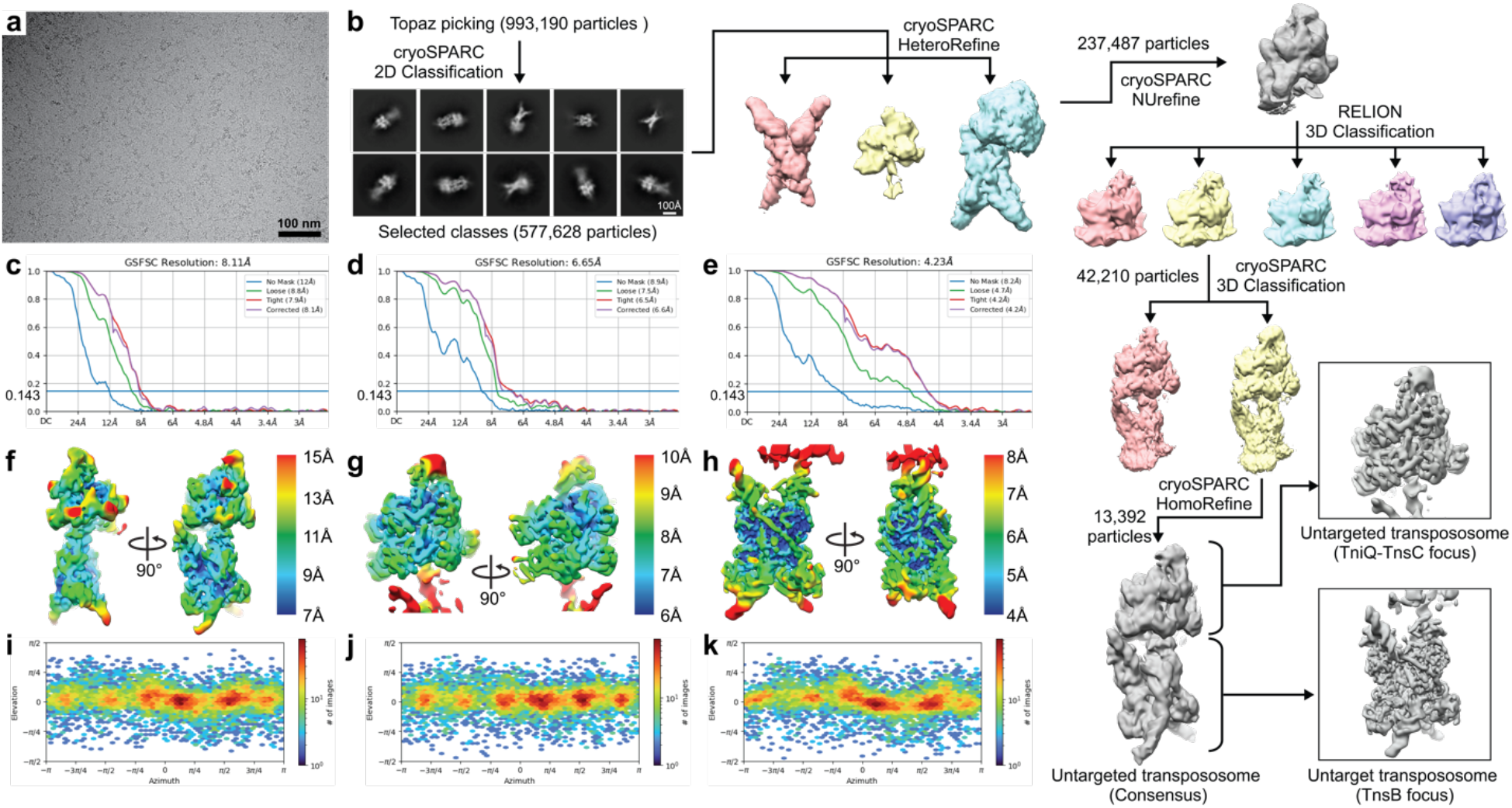
Cryo-EM image analysis of the untargeted, RNA-independent ‘BCQ’ transpososome. **a**, Representative cryo-EM micrograph of the ‘BCQ’ transpososome sample, generated by incubating TnsB, TnsC, TniQ with target-LE and target-RE DNA substrates. The black bar represents 100 nm. **b,** Computational workflow used for analyzing the ’BCQ’ transpososome cryo-EM dataset. Topaz^51^-extracted particles were first subjected to 2D classification in cryoSPARC^50^. 2D classes with cryo-EM density for the TnsB strand-transfer complex (STC) and TnsC oligomer were selected for downstream heterogeneous refinement in cryoSPARC. Particles classified as the TnsB-TnsC complex were then subjected to non-uniform refinement (NUrefine) in cryoSPARC using a mask that covers the TniQ-TnsC region. The aligned particle stacks were then further classified using 3D classification in RELION^52^, focusing on TniQ and the adjacent two subunits of TnsC. One class showing the best resolved TniQ density was selected for another round of 3D classification in cryoSPARC focusing on TniQ. The final particle stack with stronger TniQ density was then subjected to homogeneous refinement. Focused refinement was done on the TniQ-TnsC complex or the TnsB-STC region of the reconstruction. See **Methods** for details. **c-e,** Gold-standard Fourier shell correlation (GSFSC) curves from the consensus refinement (**c**) and the focused refinements of the TniQ-TnsC region (**d**) or the TnsB-STC region (**e**) of the reconstruction. The 0.143 GSFSC cutoff is indicated as a blue horizontal line. **f-h,** Estimated local resolution from the consensus refinement (**f**) and the focused refinements of the TniQ-TnsC region (**g**) or the TnsB-STC region (**h**). Local-resolution filtered reconstructions were colored based on the estimated local resolution, as indicated in the legend. **i-k,** Angular distribution heatmap of the particle stack in the final consensus refinement (**i**) and focused refinements of the TniQ-TnsC region (**j**) and the TnsB-STC region (**k**). Colors indicate particle count, as indicated in the legend.

**Supplementary Fig. 3.**
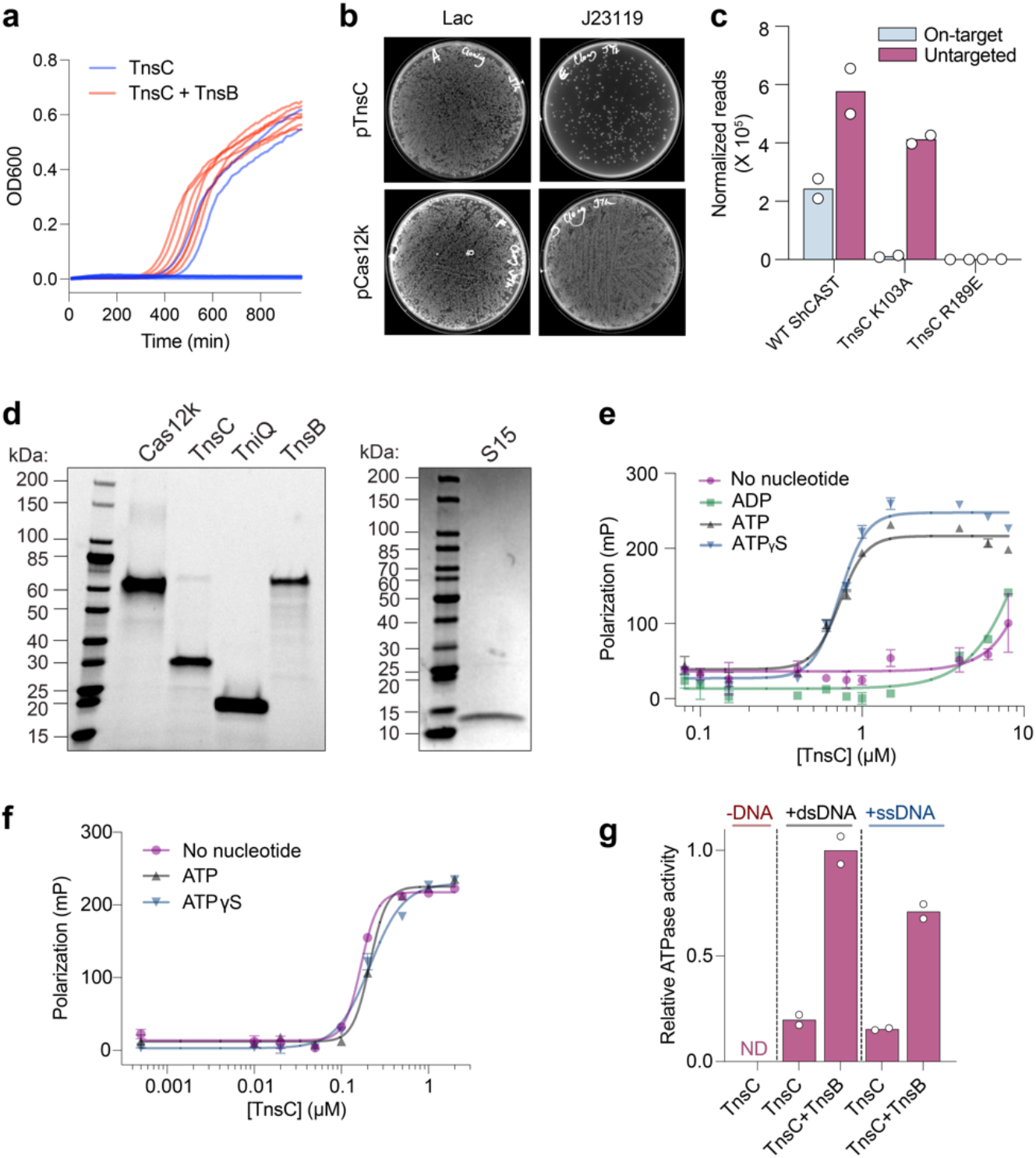
Supporting experimental data for ShCAST biochemical reconstitution. **a**, *E. coli* growth curves of seven biological replicate cultures of a TnsC over-expression strain under inducible conditions, with or without TnsB. The majority of cultures without TnsB fail to grow. **b,** Pictures of transformation plates after attempts to clone TnsC or Cas12k expression plasmids with a Lac (weak) or J23119 (strong) promoter, providing extra evidence that strong TnsC over-expression leads to cellular toxicity. The same competent cells and input DNA amounts were used in all cases, and surviving colonies with a strong promoter driving TnsC frequently exhibited mutations in the promoter or ORF. **c,** Total genome-mapping reads detected for the indicated expression construct, normalized, and scaled (**Methods**); the mean is shown from n = 2 independent biological replicates. **d,** SDS-PAGE analysis of purified ShCAST components used in biochemical transposition assays. **e,** DNA binding by TnsC is ATP dependent under 200 mM NaCl conditions, as observed using fluorescence polarization experiments with a 55-bp FAM-labeled DNA substrate. Data shown represent mean ± s.d for n = 3 technical replicates. **f,** Under low-salt conditions (50 mM NaCl), DNA binding by TnsC is nucleotide-independent. Data are shown as in **e**. **g,** TnsB stimulates the ATP hydrolysis activity of TnsC in a DNA-dependent reaction, as determined using a Malachite Green assay. The mean is shown from n = 2 technical replicates; ND, not detected.

**Supplementary Fig. 4.**
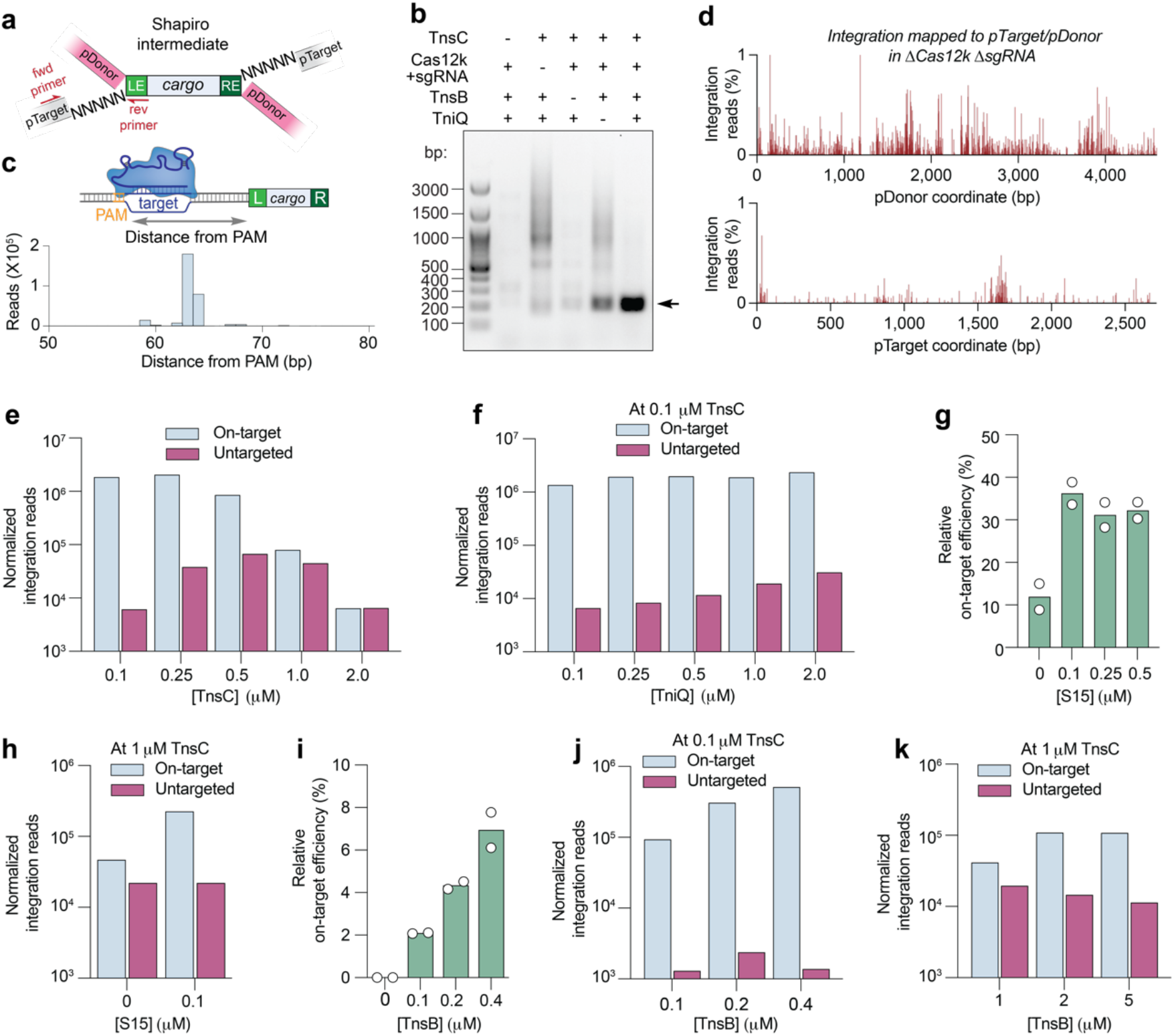
The relative stoichiometry of CAST components controls pathway choice. **a**, Biochemical transposition assays were performed *in vitro* with purified ShCAST components, followed by PCR-based amplification of the junction between pTarget and the transposon left end (LE) using the indicated primers (red). Note that ShCAST generates Shapiro intermediate products in these experiments^18^; NNNNN denotes the single-stranded target-site duplication (TSD) region. **b,** Transposition products were analyzed by PCR amplification and 1.5% agarose gel electrophoresis for the indicated conditions. The on-target product (black arrow) is only efficiently generated in the presence of all ShCAST components, though TniQ is partially dispensable for *in vitro* transposition under these conditions, as previously reported^11^. **c,** Integration site distribution for the on-target products in **b**, determined by high-throughput amplicon sequencing. **d**, Zoomed-in view of integration reads comprising 1% or less of pDonor-(top) and pTarget-(bottom) mapping reads, in a biochemical transposition assay performed without Cas12k or sgRNA. **e,** Normalized integration reads detected at on-target and untargeted sites from biochemical transposition assays, at varying TnsC concentrations. **f,** Normalized integration reads detected at on-target and untargeted sites from biochemical transposition assays, with 0.1 µM TnsC and varying TniQ concentrations. **g,** On-target transposition efficiency from biochemical transposition assays as detected by qPCR, with 0.1 µM TnsC and varying S15 concentrations. **h,** Normalized integration reads detected at on-target and untargeted sites from biochemical transposition assays with 1 µM TnsC, with or without S15. **i,** On-target transposition efficiency from biochemical transposition assays as detected by qPCR, with 0.1 µM TnsC and varying TnsB concentrations. **j,** Normalized integration reads detected at on-target and untargeted sites from biochemical transposition assays, with 0.1 µM TnsC and varying TnsB concentrations. **k**, Normalized integration reads detected at on-target and untargeted sites from biochemical transposition assays, with 1 µM TnsC and varying TnsB concentrations.

**Supplementary Fig. 5.**
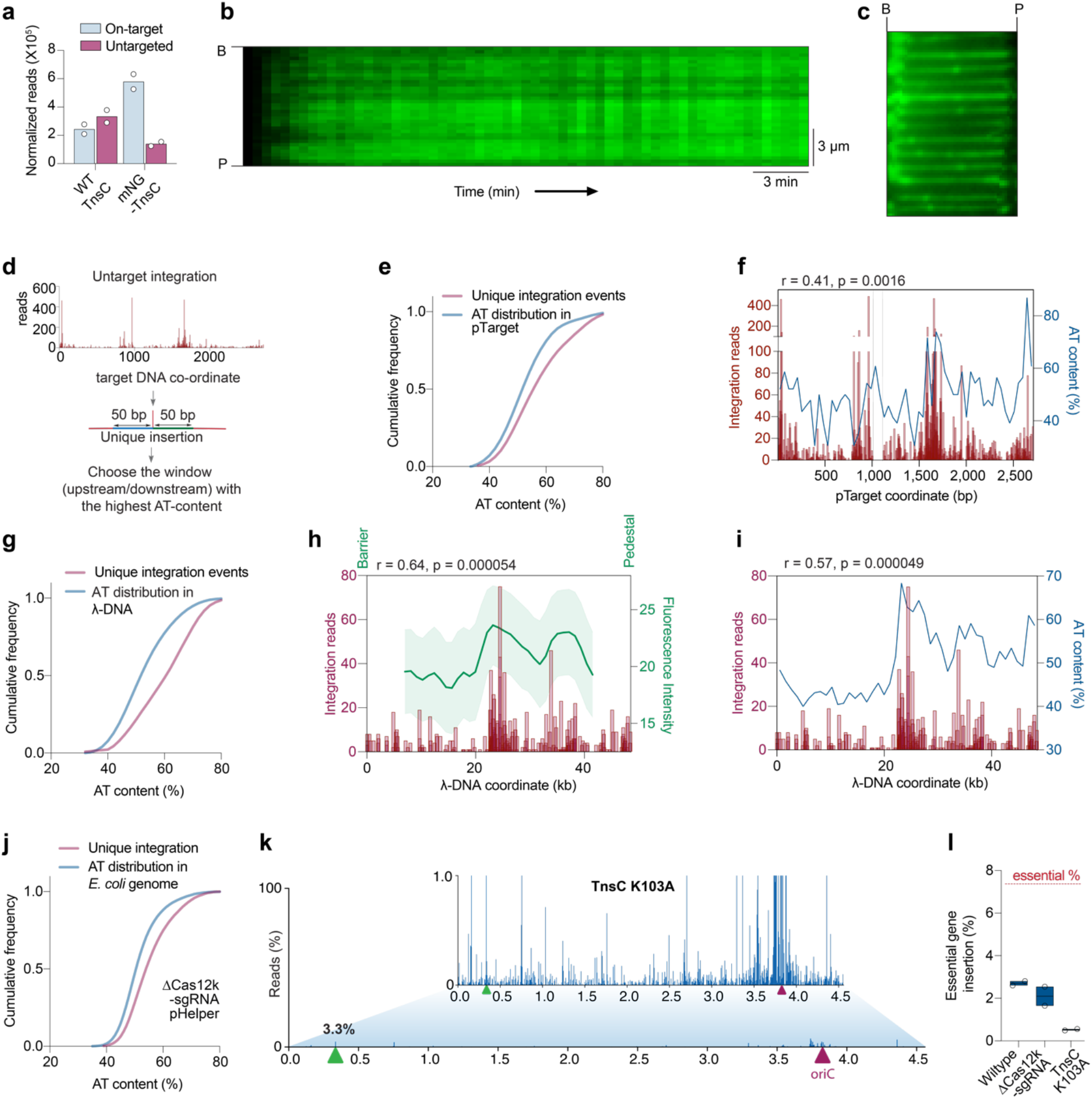
TnsC binding and TnsC-mediated integration exhibit preferential AT bias. **a**, Normalized integration reads detected at on-target and untargeted sites from cellular transposition assays, with either WT TnsC or an mNeonGreen (mNG)-TnsC fusion. **b,** Representative kymograph from a DNA curtains assay with 100 nM mNG-TnsC, showing the time-dependent evolution λ-DNA binding. B, barrier; P, pedestal. **c,** λ-DNA was completely coated with TnsC at high (500 nM) concentrations, as exemplified by a representative DNA curtains image (**Supplementary Movie 2**). **d,** Analytical workflow to investigate the AT content surrounding untargeted integration events in transposition assays. Two windows (50-bp for biochemical or 100-bp for genomic integration) flanking each unique event were analyzed, and the highest AT content was retained for subsequent analyses. A similar workflow was performed for random sampling of the same DNA substrate. **e,** Cumulative frequency distributions for the AT content within a 50-bp window flanking unique integration events on pTarget (in biochemical integration assay), using ShCAST with WT TnsC and sgRNA-6 (n = 504 unique integration events), compared to random sampling of pTarget (n = 50,000 counts; **Methods**). The distributions were significantly different, based on results of a Mann-Whitney *U* test (*P* = 3.64 x 10^-11^). **f**, Graph comparing the distribution of AT content (46-bp bins) and integration reads on pTarget, with indicated Spearman correlation coefficient and results from a two-tailed significance test. **g,** Cumulative frequency distributions for the AT content within a 50-bp window flanking unique integration events on λ-DNA, using ShCAST with WT TnsC and sgRNA-6 (n = 190 unique integration events), compared to random sampling of λ-DNA (n = 50,000 counts; **Methods**). The distributions were significantly different, based on results of a Mann-Whitney *U* test (*P* = 4.77 x 10^-11^). **h**, Graph comparing the distribution of mNG-TnsC fluorescence intensity (1,078-bp bins) and integration reads on λ-DNA, with indicated Spearman correlation coefficient and results from a two-tailed significance test. **i**, Graph comparing the distribution of AT content (1,078-bp bins) and integration reads on λ-DNA, with indicated Spearman correlation coefficient and results from a two-tailed significance test. **j,** Cumulative frequency distributions for the AT content within a 100-bp window flanking unique integration events in the *E. coli* genome, using ShCAST in the absence of Cas12k and sgRNA (n = 3,103 unique integration events), compared to random sampling of the *E. coli* genome (n = 50,000 counts; **Methods**). The distributions were significantly different, based on results of a Mann-Whitney *U* test (*P* = 4.40 X 10^-83^). **k,** Genome-wide view of *E. coli* genome-mapping reads for ShCAST with a K103A TnsC mutation; the zoomed-in view visualizes reads comprising 1% or less of genome-mapping reads. The target site is marked with a green triangle, and the *E. coli* oriC is marked with a maroon triangle. **l**, Untargeted genomic integration events are depleted in essential genes. The observed percentage of integration reads occurring within essential genes was quantified and plotted for each of the indicated datasets, from n = 2 independent biological replicates. The percentage of sequences in the *E. coli* genome denoted as essential is marked with a red line.

**Supplementary Fig. 6.**
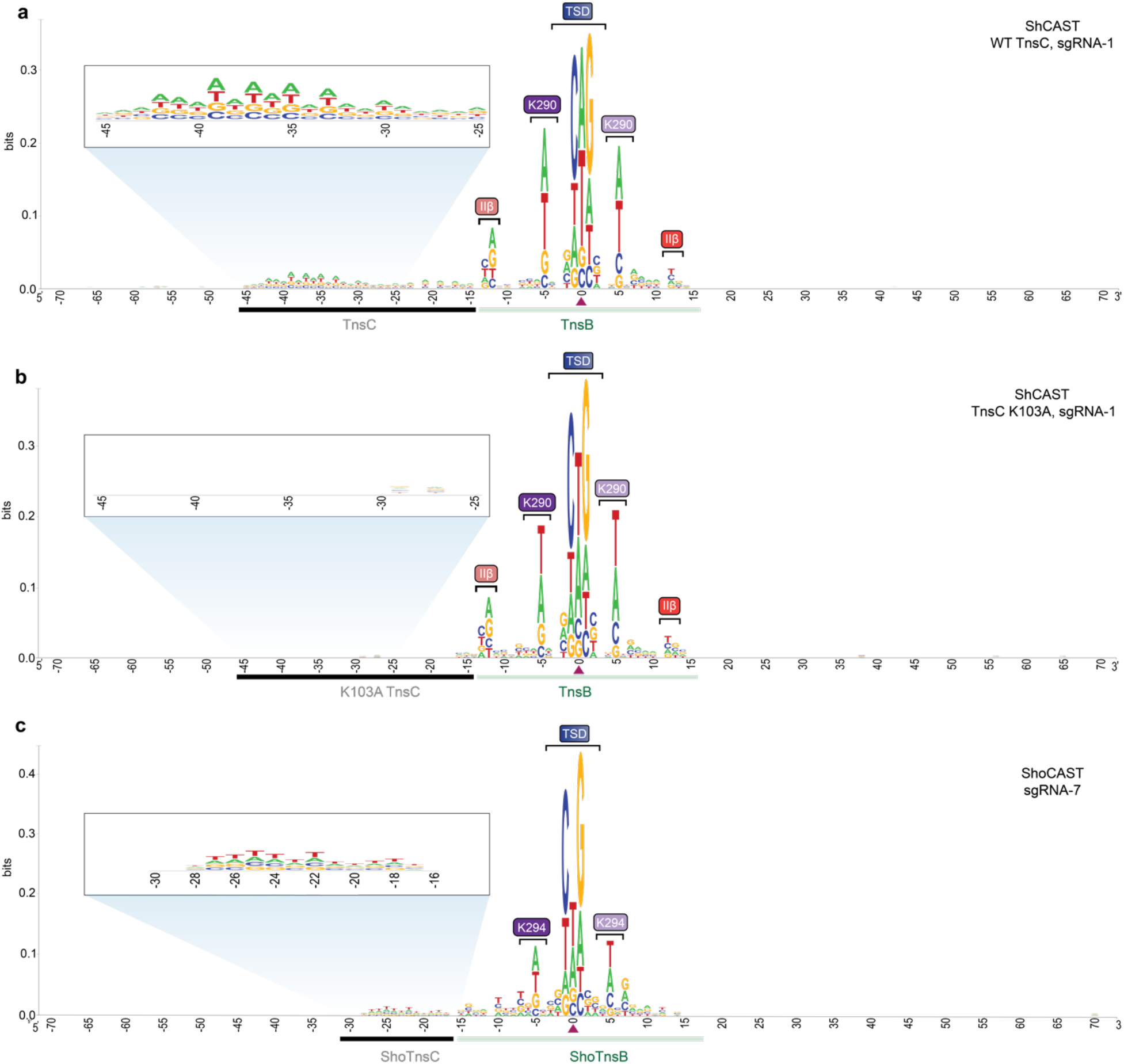
TnsC selects AT-rich features upstream of the integration site. **a**, WebLogo^60^ from a meta-analysis of untargeted genomic transposition (n = 6,800 unique integration events) from previously published data with pHelper and sgRNA-1^14^. The site of integration is noted with a maroon triangle. As with the dataset from this study (**Fig. 3h**), these results reveal an AT-rich sequence spanning ∼25 bp that likely reflects the footprint of two turns of a TnsC filament (black), next to motifs within/near the target-site duplication (TSD) region that represent TnsB-specific sequence motifs (green). Specific TnsB residues/domains contacting the indicated nucleotides are shown. The zoomed-in inset highlights periodicity in the sequence bound by TnsC. **b,** WebLogo^60^ from a meta-analysis of untargeted genomic transposition (n = 6,800 unique integration events) with a modified pHelper encoding a TnsC K103A mutation and sgRNA-1, visualized as in **a**. The AT-rich motif is conspicuously absent, as compared to experiments with WT TnsC. **c,** WebLogo^60^ from a meta-analysis of untargeted genomic transposition (n = 6,800 unique integration events) with the divergent ShoCAST system encoding sgRNA-7, visualized as in **a**. The AT-rich motif is conspicuously shorter in size with ShoCAST as compared to ShCAST, in agreement with the distinct integration distance distribution for both systems^11, 14^.

**Supplementary Fig. 7.**
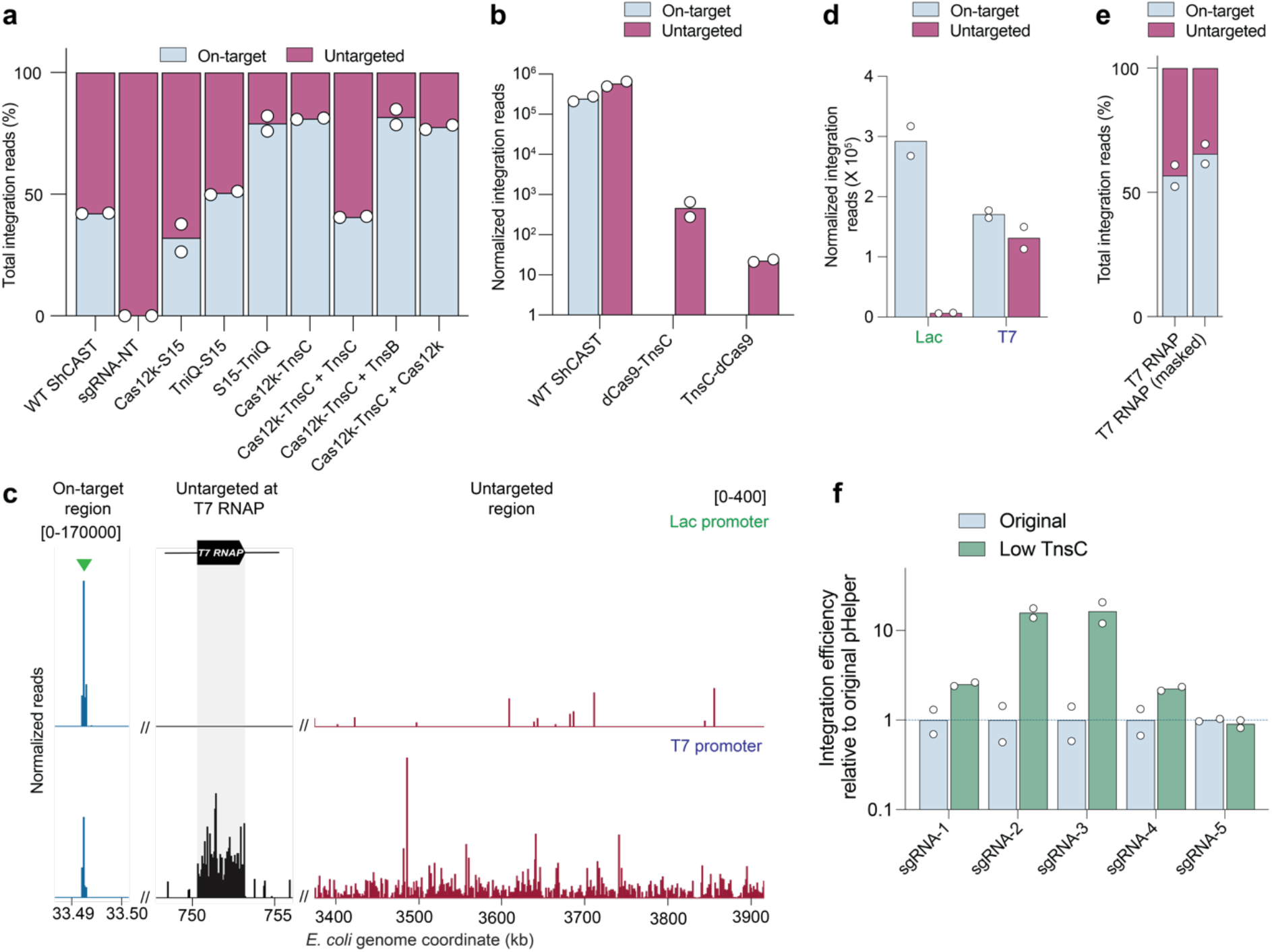
Engineering strategies to rationally alter ShCAST specificity. **a**, Fraction of total genome-mapping integration reads detected at on-target and untargeted sites, with WT ShCAST and sgRNA-1, ShCAST and a non-targeting sgRNA control (sgRNA-NT), or the listed ShCAST fusion constructs with sgRNA-1. In some cases, a fusion was supplemented with an additional plasmid encoding unfused protein components, as indicated. **b**, Normalized genome-mapping integration reads detected at on-target and untargeted sites, for WT ShCAST and N- and C-terminal dCas9-TnsC fusions. No on-target reads were obtained for these fusion constructs. **c**, Normalized integration reads detected at the on-target site (left), the T7 RNAP locus (middle), and a representative untargeted region (right), using either a TnsC expression plasmid driven by a Lac or T7 promoter. A high density of untargeted insertions was observed within the T7 RNAP gene, likely due to positive selection for loss-of-function transposition events. Note the differing y-axis ranges. **d,** Normalized genome-mapping integration reads detected at on-target and untargeted sites, when TnsC was expressed with a Lac or T7 promoter. **e**, Fraction of total genome-mapping integration reads detected at on-target and untargeted sites from experiments with T7 promoter-driven TnsC, with or without computational masking of insertions within the T7 RNAP gene. **f,** Relative on-target transposition efficiency measured for the modified design in which TnsC was expressed from a low-strength promoter compared to the original pHelper. The guide RNA are indicated sgRNA 1-5. Measurements were made by Taqman qPCR and normalized against the *rssA* locus. For **a, b, d-f**, the mean is shown from n = 2 independent biological replicates.

